# KaryoScope: rapid, alignment-free sequence annotation for the pangenome era

**DOI:** 10.64898/2026.05.15.725544

**Authors:** T. Rhyker Ranallo-Benavidez, Yi-An Chen, Tamara Potapova, Jarno Alanko, Hailey Loucks, Julian Lucas, Human Pangenome Reference Consortium (HPRC), Andrea Guarracino, Simon J. Puglisi, Camille Marchet, Karen Miga, Jennifer L. Gerton, Floris P. Barthel

## Abstract

The pangenome era is producing long-read sequencing data and complete genome assemblies (1–3) at a pace that current annotation methods cannot match. Existing tools were each built for a single feature class (repeats, centromeric satellites, or genes) and falter precisely where the genome is most variable and harbours clinically important variation: the centromeres, subtelomeres, and acrocentric short arms. Here we present KaryoScope, an alignment-free method to annotate an assembly at base-pair resolution across any desired feature classes in a single pass, completing in minutes on a standard workstation. Applied to the Human Pangenome Reference Consortium Release 2 assemblies (3), KaryoScope identifies the SST1 macrosatellite as the recurrent sequence at Robertsonian translocation fusion points (4, 5), delivers the first pangenome-wide census of D4Z4 macrosatellite structural diversity at the 4q and 10q subtelomeres relevant to facioscapulohumeral muscular dystrophy (6), and reveals previously uncharacterised centromere structural polymorphism, including chromosome-specific satellite loss and megabase-scale rearrangement validated by fluorescence *in situ* hybridization. A pre-built KaryoScope database for the human genome is distributed alongside the tool, and additional databases can be built for any reference genome or annotation source. Together, these capabilities bring the most variable regions of the genome within reach for comparative, clinical, and pangenome-scale analysis. KaryoScope is available at https://github.com/barthel-lab/KaryoScope.

## Introduction

The pangenome era has shifted the bottleneck in genomics from sequence acquisition to sequence interpretation. Long-read sequencing now routinely yields complete assemblies, once produced only by dedicated consortia (1–3) but increasingly delivered by individual laboratories (7–10) and across population-scale collections (11). Hundreds of phased diploid haplotypes are now available, yet no existing annotation tool delivers a unified, genome-wide, base-pair resolution interpretation of these assemblies. Available tools each handle only a single feature class, and they perform worst in the satellite-dense centromeres, subtelomeres, and short arms of the acrocentric chromosomes, regions that underlie clinically important rearrangements such as Robertsonian translocations (4, 5) and facioscapulohumeral muscular dystrophy type 1 (FSHD1) (6).

Annotation has historically been split across tools specific to each feature class: RepeatMasker (12) and curated libraries (Dfam (13), Repbase (14)) for interspersed elements; CenSat, HumAS-HMMER (15), CenMAP (16), and Centeny (17) for centromeric satellites; and Liftoff (18) and BRAKER (19) for genes. Each performs well within its remit, but none integrates these feature classes into a single annotation, and none scales to the throughput pangenome assemblies now demand.

Two practical consequences follow. First, satellite-dense regions are precisely where alignment-based pipelines perform worst: repetitive sequence, reference bias, and inter-individual structural variation conspire to produce ambiguous alignments and systematic miscalls. Second, existing pipelines were built for one genome at a time, taking hours to days per haplotype on dedicated hardware, incompatible with pangenome cohorts of hundreds of phased assemblies. Pangenome-scale annotation therefore requires a method that scales to minute-per-haplotype runtimes and annotates multiple features in tandem.

Here, we introduce KaryoScope, an alignment-free annotation tool that assigns each *k*-mer in a query assembly to a feature drawn from one or more user-defined hierarchical feature sets, producing a base-pair resolution annotation in a single pass. Because a feature set is simply any tiling of a reference with labelled regions, KaryoScope is extensible to arbitrary annotation sources, from satellite catalogs and repeat libraries to cytobands, FISH-probe coordinates, structural-variant break-points, and metagenomic taxonomies. We distribute a pre-built database for the human genome, derived from T2T-CHM13v2.0 (1), with six feature sets covering chromosome of origin, satellite composition, interspersed repeats, sub-telomeric structure, gene boundaries, and acrocentric-specific features; additional databases can be built for any reference or community-curated annotation. We have previously applied KaryoScope to fibroblast cell-line assemblies (7) and to the HG008 tumour-normal pair (10), and a performance-optimized variable-length *k*-mer index, HKS (20, 21), provides an improved query backend at pangenome scale.

We apply KaryoScope to the HPRC Release 2 pangenome (3) across the satellite-dense regions where annotation tools have, until now, fallen short. We first establish the method on the HG002 diploid assembly and benchmark its annotations against RepeatMasker and CenSat. We then present three applications enabled by pangenome-scale annotation. First, KaryoScope identifies the SST1 macrosatellite as the recurrent sequence at the fusion points of three Robertsonian translocation cell lines, resolving the mechanism of acrocentric fusion from *k*-mer composition alone. Second, KaryoScope delivers the first pangenome-wide census of D4Z4 macrosatellite structural diversity at the 4q and 10q subtelomeres, including non-canonical configurations relevant to FSHD1. Third, KaryoScope reveals previously uncharacterized centromere structural polymorphism, with selected variants on chromosomes 3, 5, and 9 validated by fluorescence *in situ* hybridization.

By enabling rapid, alignment-free, base-pair resolution annotation across any user-defined feature classes, KaryoScope brings the most variable regions of the genome within reach for comparative, clinical, and pangenome-scale analysis. The same architecture applies wherever a reference annotation exists, from satellite catalogs in the human genome to gene and repeat libraries in other species, positioning KaryoScope as a general framework for annotating complete assemblies as they become available across the tree of life.

## Results

### A single KaryoScope index produces a base-pair resolution diploid annotation of HG002

To demonstrate KaryoScope’s ability to produce a comprehensive, annotation of a genome assembly, we annotated the HG002 diploid assembly (NA24385, Ashkenazi, 46,XY) from the Human Pangenome Reference Consortium (2) using a *k*-mer database constructed from the T2T-CHM13v2.0 reference (1). The database comprises six hierarchically organized feature sets: chromosome, region, repeat, gene, acrocentric, and subtelomeric, each providing a distinct layer of annotation for the same underlying *k*-mers (see Methods).

KaryoScope uses these annotation layers to produce a karyotype-style summary of the assembly. KaryoScope uses the chromosome feature set to assign each contig to a chromosome of origin based on majority k-mer content. It then uses the region feature set (p arm, centromere, q arm) to orient each contig and order multi-piece chromosomes. Annotations are summarized at user-configurable resolutions (e.g., 1 Mb, 100 kb, 100 bp) for visualization at scales ranging from genome-wide (Figure 1; Supplementary Figure S1) to per-region, including centromeric (Supplementary Figure S2) and subtelomeric (Supplementary Figure S3 and Supplementary Figure S4) views.

**Fig. 1.**
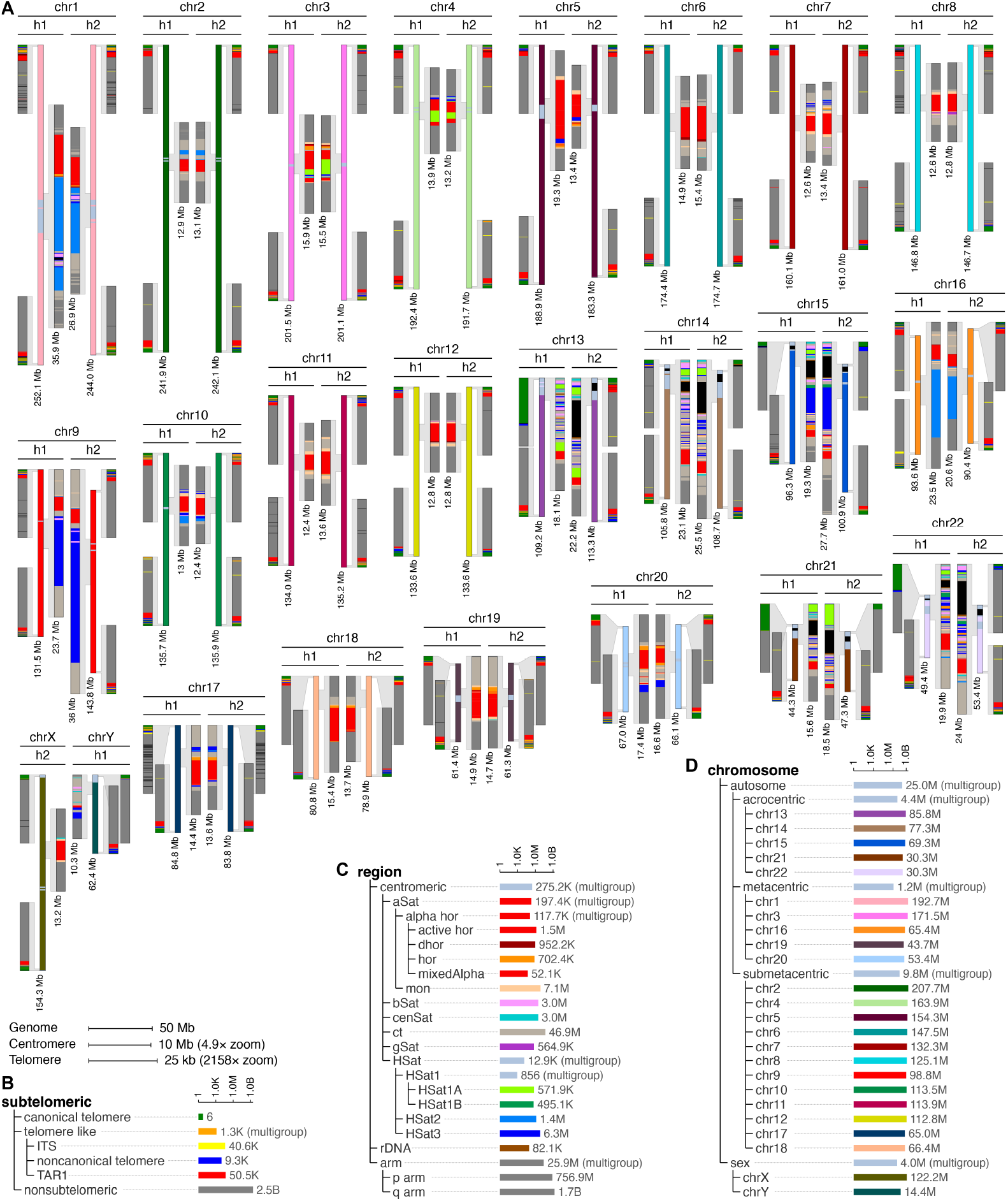
KaryoScope karyotype of the HG002 diploid assembly with centromere and subtelomere zoom views. **(A)** All 24 chromosomes are displayed: both haplotypes (h1, h2) for the 22 autosomes are shown side by side, mirror-imaged about the center, with chrX and chrY shown next to each other. Reading outward from the mirror axis, each haplotype comprises a centromere zoom panel (100 kb bins) showing satellite composition from the region feature set (HOR subfamilies including active, inactive, and divergent HORs; mixed and monomeric *α*-satellite; *β* -satellite; cenSat; centromeric transition (ct) regions; gSat; human satellite subfamilies including HSat1A, HSat1B, HSat2, and HSat3; rDNA), a full chromosome track (1 Mb bins) coloured by chromosome of origin, and a subtelomere zoom panel (100 bp bins) showing subtelomeric features (canonical telomere, noncanonical telomere, ITS, TAR1). This arrangement places the centromere zooms adjacent for direct haplotype-to-haplotype comparison. Positions whose *k*-mers are absent from the database (“novel”) are coloured black; the prominent black blocks on the short arms of the acrocentric chromosomes correspond to assembly gaps (stretches of Ns) within the rDNA arrays, which are not yet fully resolved in the HG002 assembly. Scale bars: genome, 50 Mb; centromere, 10 Mb (4.9× zoom); subtelomere, 25 kb (2,158× zoom). **(B)** Subtelomeric feature set hierarchy tree with log-scaled *k*-mer count bars. **(C)** Region (satellite) feature set hierarchy tree. **(D)** Chromosome feature set hierarchy tree. Assembly: HG002 (NA24385, Ashkenazi, 46,XY) from the Human Pangenome Reference Consortium.

KaryoScope correctly assigned chromosome of origin across both haplotypes for all 46 chromosomes of HG002 (Figure 1A; Supplementary Figure S1B). At the genome scale (1 Mb bins), chromosome of origin annotations were uniform across each chromosome arm, with centromeric regions clearly delineated by the transition to satellite annotations in the region feature set (Supplementary Figure S2A).

Centromeric satellite composition varied substantially across chromosomes (Figure 1A; Supplementary Figure S2A). Chromosomes 1 and 16 harbored large HSat2 arrays flanking their *α*-satellite cores, while the acrocentric chromosomes (chr13–15, chr21–22) showed characteristic rDNA-adjacent centromeric organization. Haplotype-specific differences in satellite array length and composition were visible at multiple chromosomes, consistent with the known rapid evolution of centromeric repeats (15).

At the subtelomeric scale (100 bp bins), canonical (TTAGGG)_*n*_ telomere tracts were present at all chromosome termini in both haplotypes, with tract lengths varying from ∼2 kb to *>*15 kb (Figure 1A; Supplementary Figure S3A). TAR1 elements were identified at most chromosome ends. The region feature set additionally revealed *β* -satellite arrays at the 4q and 10q subtelomeres (Supplementary Figure S4A), marking the D4Z4 macrosatellite arrays implicated in FSHD1 (22).

### KaryoScope matches RepeatMasker and CenSat at *>*95% base-pair accuracy

To evaluate annotation accuracy, we compared KaryoScope to the two community standards for repeat (RepeatMasker (12)) and centromeric-satellite (CenSat (15)) annotation. We performed a self-benchmark on T2T-CHM13v2.0 (from which the KaryoScope database was constructed) and a cross-genome benchmark on the HG002 diploid assembly using HG002’s own published annotations as truth (Supplementary Note 1). Because RepeatMasker collapses Satellite, Simple_repeat, and Low_complexity into related *k*-mer pools that cannot be cleanly distinguished at the *k*-mer level, these classes were merged into a single tandem-repeat category for evaluation; less-specific calls within the feature hierarchy were classified as *compatible* rather than incorrect (see Methods).

KaryoScope’s repeat annotations matched RepeatMasker at 98.7% of *k*-mer positions on T2T-CHM13v2.0 and at 95.2– 95.3% on the two HG002 haplotypes, with an additional ∼1% compatible at a less specific hierarchy level (Figure 2A, C; Supplementary Figure S5A, C). The major interspersed repeat classes each matched at *>*94% on HG002 (LINE 98.3%, LTR 97.7%, DNA 96.8%, SINE 94.7%), with cross-genome differences from the self-benchmark attributable to genuine inter-individual variation in transposable element content rather than misclassification. Hierarchy-aware smoothing was central to this accuracy: prior to smoothing, ∼85% of HG002 positions matched exactly and ∼9% were compatible; after smoothing, nearly all compatible calls were resolved to their most specific labels.

**Fig. 2.**
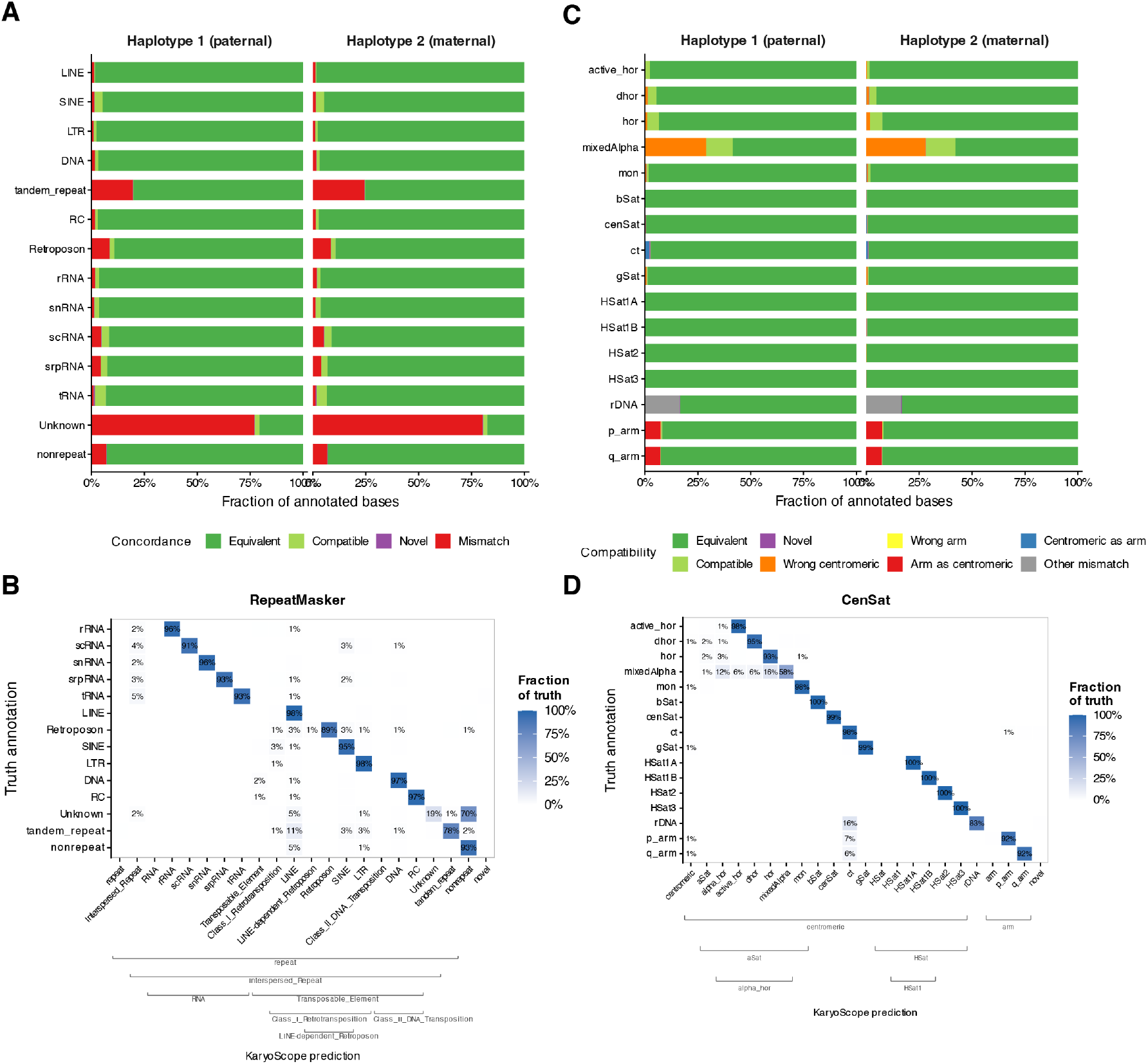
KaryoScope concordance with RepeatMasker and CenSat on the HG002 diploid assembly. KaryoScope was applied to the HG002 diploid assembly (haplotype 1, paternal; haplotype 2, maternal) using the T2T-CHM13v2.0-derived *k*-mer database, and the resulting annotations were compared to HG002’s own RepeatMasker and CenSat v2.0 annotations as truth. **(A)** Per-category concordance for the repeat feature set (RepeatMasker truth) across both haplotypes. Bars show the fraction of annotated bases per category that match the truth annotation exactly (Equivalent), at a less specific level in the feature hierarchy (Compatible), as a previously unannotated repeat (Novel), or as an incorrect class (Mismatch). **(B)** Per-category concordance for the region (centromeric satellite) feature set (CenSat truth) across both haplotypes, with mismatches further partitioned into wrong centromeric subtype, wrong arm, arm-as-centromeric, centromeric-as-arm, and other. **(C)** Confusion matrix showing KaryoScope predictions (columns) versus RepeatMasker truth (rows) collapsed across both haplotypes. Cell shading encodes the fraction of each truth category assigned to each predicted category. Brackets along the bottom axis indicate the hierarchical structure of the repeat feature set, allowing compatible parent-category predictions to be distinguished from outright mismatches. **(D)** Confusion matrix for the region feature set against CenSat truth, with hierarchical brackets for centromeric and arm subcategories. Assembly gaps were excluded from the evaluation in all panels.

Centromeric-satellite annotations matched CenSat at 99.4% on T2T-CHM13v2.0 and at 98.2–98.7% within centromeric and rDNA regions of HG002, with per-subtype accuracy of 99.8–100% across HSat1A, HSat1B, HSat2, HSat3, and the principal *α*-satellite HOR families (Figure 2B, D; Supplementary Figure S5B, D). Genome-wide concordance against CenSat was lower (92.7–93.3%); the dominant source of discordance was a 6.6–6.8% population of arm positions annotated as the centromere-transition (ct) label, virtually identical across T2T-CHM13v2.0 and both HG002 haplotypes. The ct label in CenSat v2.1 marks segmental duplications interspersed between centromeric satellite arrays, and KaryoScope’s recognition of paralogous *k*-mer content between these regions and the chromosome arms reflects genuine sequence identity rather than a smoothing artifact (Supplementary Note 1). A single, reference-derived *k*-mer database therefore reproduces both community-standard annotations at *>*95% base-pair accuracy across the entire diploid genome, including its most repetitive compartments.

The released KaryoScope pipeline (KaryoScope-KMC) annotates a diploid human assembly across chromosome, region, repeat, gene, acrocentric, and subtelomere feature sets in a single pass in 7.7 ± 0.3 minutes per haplotype on 16 CPU threads with under 50 GiB of peak memory (*n* = 3; Intel Xeon Ice Lake, 2.80 GHz; Supplementary Table S1; Supplementary Note 2), approximately 100× faster than Repeat-Masker on equivalent hardware. The two principal steps, *k*-mer feature assignment and hierarchy-aware smoothing, are independently parallelizable. Replacing the KMC query backend with the Spectral Burrows–Wheeler Transform-based HKS index (20, 21) and porting the smoothing step to Rust (KaryoScope-HKS) further reduces end-to-end per-haplotype runtime to ∼2 minutes and peak memory to under 11 GiB on the same hardware (Supplementary Figure S6; Supplementary Note 2), more than 300× faster than RepeatMasker.

### KaryoScope localizes Robertsonian translocation breakpoints to the SST1 macrosatellite

Formed by fusion of the long arms of two acrocentric chromosomes, Robertsonian translocations (ROBs) are the most common structural rearrangement in humans (approximately 1 in 1,000 live births) and a recurrent cause of reproductive loss and meiotic aneuploidy. Classical molecular studies localized ROB breakpoints to the pericentromeric satellite regions of the acrocentric short arms but left the mechanism and the precise recombination substrate unresolved (23). Pangenome analyses of the acrocentric short arms subsequently resolved the SST1 macrosatellite, an array of tandem ∼1.4 kb repeats shared between chromosomes 13, 14 and 21, as a candidate recombination substrate for ROB formation (4), a mechanism confirmed directly by complete assembly of three ROB-carrying cell lines and identification of partial SST1 monomers with diagnostic sequence asymmetry flanking each fusion (5). To ask whether KaryoScope’s *k*-mer composition alone is sufficient to recover this mechanism without requiring alignment, we re-analysed the three published Verkko assemblies (GM03417, rob(14;21)(q10;q10); GM03786, rob(13;14)(q10;q10); GM04890, rob(13;14)(q10;q10)).

KaryoScope independently identified all three fusion scaffolds as contigs carrying contiguous chromosome-specific *k*-mer signatures from two distinct acrocentrics (Figure 3A–C). Multi-level zoom visualization spanning 20 Mb down to 200 kb revealed, in each sample, a sharp transition between the two fusion partners’ *k*-mer content, with the SST1 satellite block (visible in the acrocentric track) flagged at the fusion boundary. SST1 block sizes varied widely across samples (69.5 kb in GM03417, 10.3 kb in GM03786, 60.9 kb in GM04890), consistent with the per-chromosome SST1 tract-length polymorphism previously reported in ROB carriers. Each main SST1 array (corresponding to the previously described SST1 subfamily 1 array) was flanked by two smaller partial arrays (∼ 0.8–1.9 kb; corresponding to the SST1 type 2L and type 2S monomers, derived from chr13/chr21 and chr14, respectively), and examination of the chromosome-specific *k*-mer content across all three array types revealed a clear pattern: the flanking partial arrays carried *k*-mers predominantly from a single fusion partner, consistent with their position on either side of the fusion, while the main arrays contained *k*-mers from both fusion partners (Supplementary Figure S7D). This *k*-mer-based identification of flanking partial SST1 monomers carrying single-partner *k*-mer content independently reproduces the breakpoint signature previously identified through sequence comparison to T2T-CHM13v2.0.

**Fig. 3.**
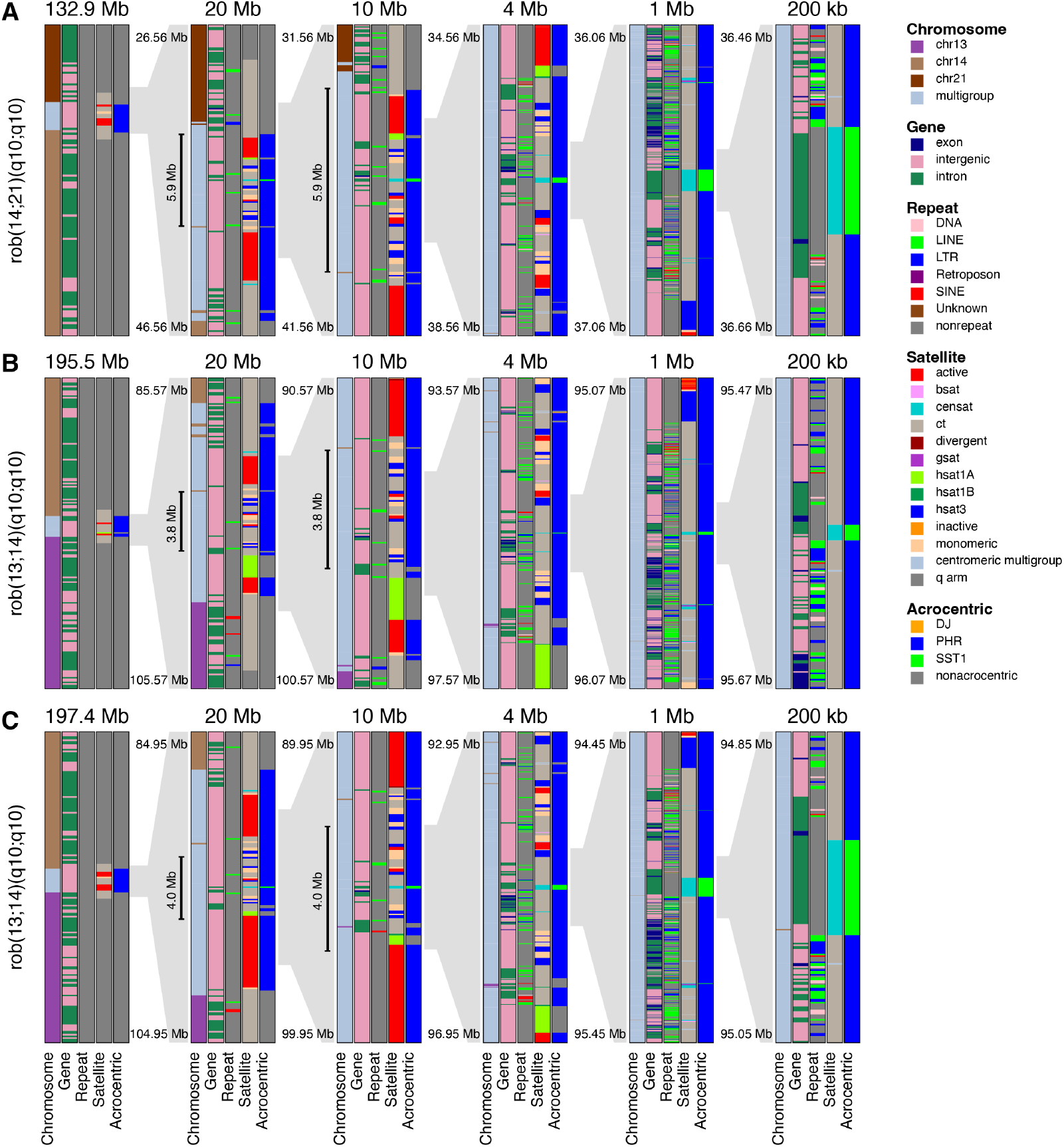
KaryoScope resolves Robertsonian translocation breakpoints at SST1 satellite repeats. Three Robertsonian translocation cell lines are shown in rows, each with six progressively zooming panels (full contig to 200 kb) centered on the SST1 satellite block at the fusion site. Five annotation tracks are displayed per panel: chromosome of origin (one colour per chromosome), gene boundaries (exon, intron, intergenic), repeat classification (DNA, LINE, LTR, Retroposon, SINE, Unknown), satellite subtypes (HOR subfamilies including active, inactive, and divergent HORs; monomeric *α*-satellite; *β* -satellite; cenSat; centromeric transition (ct) regions; gSat; human satellite subfamilies including HSat1A, HSat1B, and HSat3), and acrocentric-specific features (distal junction (DJ), pseudohomologous region (PHR), SST1 macrosatellite). **Top row:** GM03417, 45,XX,rob(14;21)(q10;q10): chr21q fused to chr14q on a 132.9 Mb contig with a 69.5 kb SST1 block at the fusion site. **Middle row:** GM03786, 45,XX,rob(13;14)(q10;q10): chr14q fused to chr13q on a 195.5 Mb contig with a 10.3 kb SST1 block. **Bottom row:** GM04890, 45,XX,rob(13;14)(q10;q10): chr14q fused to chr13q on a 197.4 Mb contig with a 60.9 kb SST1 block. All three breakpoints coincide with SST1 satellite repeat blocks (green, Acrocentric track), though SST1 block sizes vary considerably across samples.

**Fig. 4.**
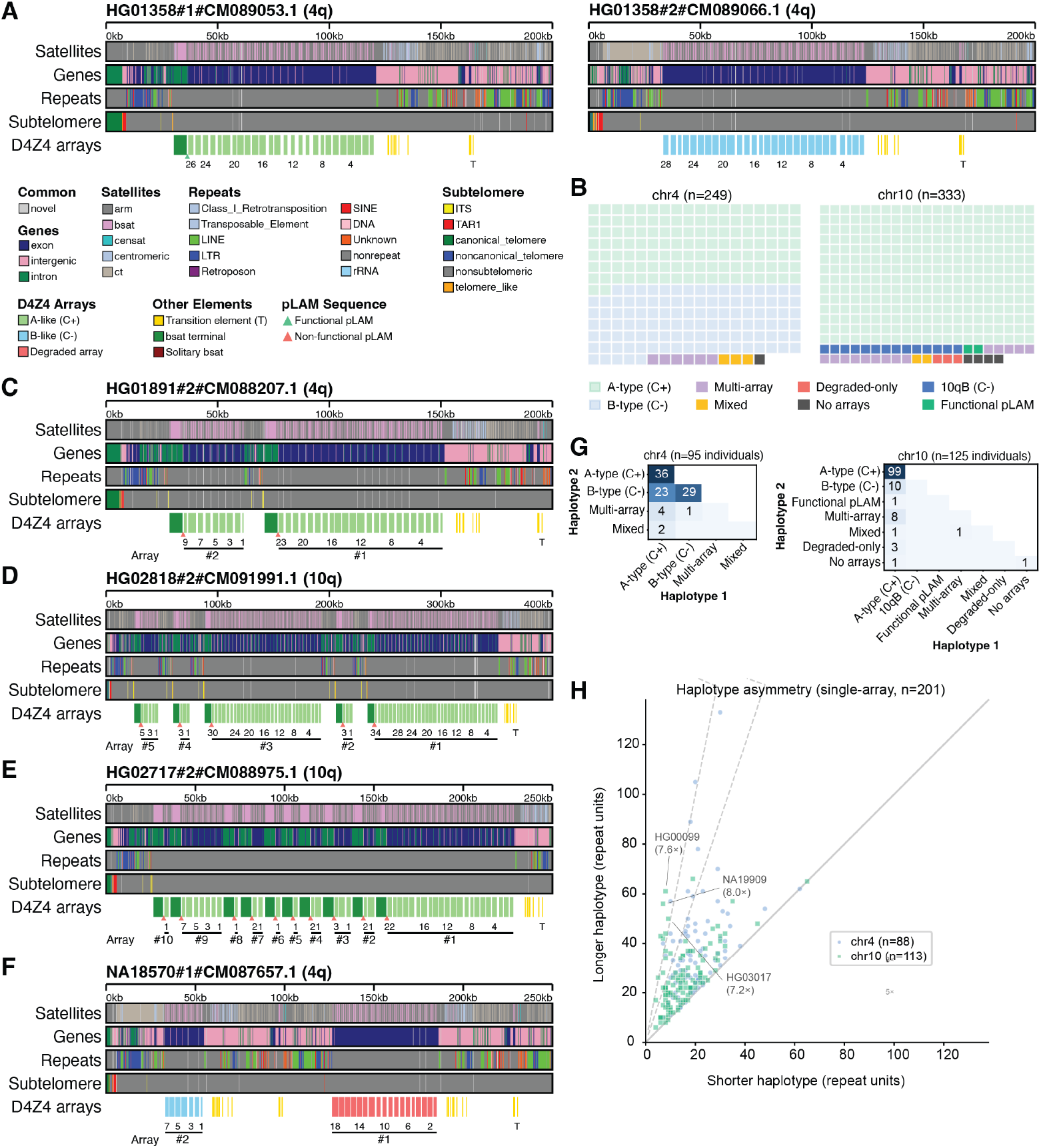
KaryoScope resolves D4Z4 macrosatellite structural variation at 4q and 10q subtelomeres. **(A)** Representative KaryoScope multi-track visualization of sample HG01358 showing both chr4 haplotypes with satellite, gene, repeat, and subtelomeric annotations. D4Z4 arrays are classified as A-type (C+, *β* -satellite terminal) or B-type (C-, segmental-duplication enriched terminal) with individually numbered repeat units. C+ denotes the presence of a *β* -satellite cap; C-denotes its absence. **(B)** Waffle charts showing D4Z4 structural configuration frequencies for chr4 (*n* = 249 haplotypes) and chr10 (*n* = 333 haplotypes) from HPRC assemblies. Categories: A-type (C+) for haplotypes with a *β* -satellite terminal and without functional pLAM, B-type (C-) for chr4 haplotypes with a segmental-duplication enriched terminal, 10qB (C-) for chr10 haplotypes with a segmental-duplication enriched terminal, Functional pLAM for haplotypes with a *β*-satellite terminal and a functional ATTAAA polyadenylation signal, multi-array, mixed, degraded-only, and no arrays. A-type (C+) and Functional pLAM together comprise all A-type haplotypes (296 on chr10, of which 2 carry functional pLAM; 138 on chr4, of which 63 carry functional pLAM). **(C–E)** Multi-array haplotypes carrying two or more distinct D4Z4 tracts: HG01891 (chr4), HG02818 (chr10), and HG02717 (chr10). **(F)** Mixed configuration in which a terminal-bearing canonical array coexists with a more proximal array lacking any detectable terminal: NA18570 (chr4). **(G)** Contingency tables of diploid configuration combinations for chr4 (*n* = 95 individuals) and chr10 (*n* = 125 individuals). Tables include only individuals for whom both haplotypes of the corresponding chromosome are telomere-to-telomere. **(H)** Haplotype asymmetry scatter plot comparing repeat unit counts between haplotypes for single-array individuals (*n* = 201), with fold-difference lines (3×, 5×) and labelled outliers.

In addition to the expected acrocentric chromosome-specific *k*-mers (chr13, chr14, chr21), the main SST1 arrays contained small numbers of *k*-mers specific to non-acrocentric chromosomes, including chr7, chr9, chr17, and chr20 (Supplementary Figure S7D). In T2T-CHM13v2.0, each of these 31-mers fell within a pericentromeric region on its respective chromosome (Supplementary Figure S7E), precisely where dispersed SST1 monomers and duplications have been annotated in the T2T-CHM13v2.0 assembly (24), and consistent with earlier reports of SST1-related sequences on chr9 (25) and an SST1 element adjacent to the HSat3 array on chr17 (1). The presence of these non-acrocentric *k*-mers within the ROB SST1 arrays indicates that SST1 repeat units on acrocentric and non-acrocentric chromosomes share substantial sequence identity at the 31-mer scale, suggesting ongoing or ancestral sequence exchange between pericentromeric SST1 elements across the genome. Whether this reflects a broader potential for SST1-mediated interchromosomal recombination beyond the acrocentric chromosomes remains to be determined, but it is notable that KaryoScope’s *k*-mer annotation recovers this cross-chromosomal homology without explicit sequence alignment.

To obtain a quantitative breakpoint interval in an alignment-free manner, we developed a symmetric cumulative *k*-mer analysis at 100 bp resolution (see Methods), inferring the breakpoint as the position along each fusion scaffold that maximally separates the chromosome-specific *k*-mer content of the two fusion partners. Applied to the three ROB assemblies, the inferred breakpoint fell within approximately 200–525 kb of the SST1 block midpoint in every case (Supplementary Figure S7A–C). This provides *k*-mer-based, alignment-free corroboration that SST1 is the recombination substrate for Robertsonian fusion.

Our analysis is complementary to the prior breakpoint localization, which used alignment to T2T-CHM13v2.0 to place each breakpoint within the SST1 array by identifying partial SST1 monomers flanking the fusion that carried diagnostic sequence differences between the two fusion partners.

The two approaches exploit the same underlying evidence, chromosome-specific sequence flanking the fusion, and converge on the same conclusion, but differ in methodology: alignment-based localization requires a contiguous assembly aligned to a reference and a prior model identifying SST1 as the substrate, whereas cumulative *k*-mer analysis requires only the pre-computed KaryoScope database and is agnostic to the identity of the recombination substrate. This distinction is practical rather than fundamental, both ultimately derive from the same reference genome, but the *k*-mer approach avoids the computational cost and complexity of alignment in repetitive pericentromeric regions, and can be applied as a rapid screen to any new assembly without bespoke analysis.

### D4Z4 macrosatellite diversity extends beyond canonical 4qA/4qB across the HPRC pangenome

The D4Z4 macrosatellite is a particularly instructive target for pangenome-scale KaryoScope annotation. Near-identical arrays of ∼3.3 kb repeat units occur on the 4q35 and 10q26 subtelomeres, and the two loci share extensive sequence identity. The structural configurations of these arrays, canonical A or B terminal feature, array length, internal integrity, and inter-array spacing, determines disease risk in FSHD1 (6). Short-read and targeted sequencing approaches routinely confound the two loci and struggle to resolve array length, terminal identity, and the full spectrum of non-canonical arrangements. HPRC Release 2 provides, for the first time, a sample size sufficient to catalog D4Z4 structural variation across hundreds of phased diploid haplotypes.

We applied KaryoScope to diploid HPRC Release 2 assemblies (3) and developed a classification pipeline (see Methods) that reconstructs D4Z4 arrays from the 4q and 10q subtelomeres and classifies each array by its distal terminal feature: a terminal *β* -satellite element (a structural proxy for the 4qA allele) or a segmental-duplication enriched region (marking the 4qB allele) (Figure 4A; Supplementary Figure S8A). Repeat-unit counts derived from KaryoScope were concordant with orthogonal HMM-based counts (Pearson *r* = 0.998; Supplementary Figure S8B), demonstrating that *k*-mer-based array reconstruction recovers unit counts at fidelity comparable to dedicated HMM-based satellite counting.

Across 249 T2T 4q haplotypes and 333 T2T 10q haplotypes from HPRC Release 2, the majority of D4Z4 arrays fell into the two canonical classes (A-type and B-type). On chromosome 4, 53.8% of haplotypes carried the A-type (*β* -satellite terminal) configuration and 45.8% the B-type (segmental-duplication enriched terminal); chromosome 10 was predominantly A-type (93.7%) with a small minority of B-type haplotypes (4.2%) (Figure 4B). The remaining haplotypes on both chromosomes fell into non-canonical configurations described below. A BLAST-based search for the canonical pLAM polyadenylation sequence within each chromosome 4 A-type BSR region identified a functional ATTAAA signal in 63 of 138 haplotypes (45.7%); the remainder carried the non-functional ATCAAA variant.

Beyond the canonical A and B configurations, KaryoScope resolved a spectrum of non-canonical architectures that have, until now, been difficult to enumerate systematically, both because such configurations are individually rare and because alignment-based approaches struggle to anchor D4Z4 events to specific 4q or 10q assembly coordinates. Notably, functional ATTAAA pLAM sequences were detected on a small number of chromosome 10 A-type arrays (Supplementary Figure S9A), departing from the 4q-specific distribution of the functional signal assumed in most targeted assays. A small number of haplotypes carried only degraded arrays in which no canonical terminal was detected, and rare haplotypes on both chromosomes contained no detectable D4Z4 array within the 250 kb subtelomeric window (Supplementary Figure S9B, C). Multi-array haplotypes carrying two or more distinct D4Z4 tracts were detected on both chromosomes, with inter-array spans dominated by novel sequence, intergenic stretches, and interspersed repeats (Figure 4C, D, E; Supplementary Figure S8C; Supplementary Figure S9D–F). Mixed arrangements, in which a terminal-bearing canonical array coexisted with a more proximal array lacking any detectable terminal, were observed on both chromosomes (Figure 4F). Diploid contingency tables show the full range of canonical pairings across the pangenome: A/A, A/B, and B/B combinations on chromosome 4, and A/A and A/B combinations on chromosome 10 (Figure 4G). Comparison of repeat-unit counts between haplotypes within single-array individuals revealed substantial intra-individual asymmetry, with several samples exceeding 3-fold differences and rare outliers surpassing 5-fold (Figure 4H; Supplementary Figure S8D). Together, these observations establish that D4Z4 structural diversity extends well beyond the canonical A/B dichotomy and, by anchoring each configura-tion to phased assembly coordinates, place this diversity in pangenome-scale context for the first time.

### KaryoScope reveals centromeric structural polymorphism across the HPRC pangenome

Centromeres are the most repeat-dense regions of the human genome, composed of megabase-scale tandem arrays of *α*-satellite HORs flanked by HSat families and *β* -satellite elements (15). These regions direct kinetochore assembly and chromosome segregation, and their sequence organization underlies the chromosome-specific and subfamily-specific identity of every human centromere (17). Centromeres are also among the most rapidly evolving regions of the human genome, exhibiting at last a 4.1-fold elevation in single-nucleotide variation over their unique flanks and up to three-fold variation in *α*-satellite HOR array length between individuals (11). Recent, pagenome-scale efforts have begun to expose the magnitude of this variation, cataloging hundreds of previously undescribed centromere haplotypes and thousands of novel HOR variants across globally diverse individuals (16). Existing annotation approaches, however, were not designed to capture this variation at pangenome scale. Consensus-matching pipelines such as RepeatMasker collapse related satellite subfamilies into generic calls, while HMM-based pipelines such as HumAS-HMMER and CenSat resolve HOR subfamilies at monomer resolution but are restricted to centromeric and pericentromeric regions. Satellite-aware annotation that scales across hundreds of haplotypes is therefore a prerequisite for any systematic survey of centromere structural variation.

To survey centromere variation at pangenome scale, we applied KaryoScope to the HPRC Release 2 assemblies (3). Each centromere was first assessed with NucFlag, an assembly-error detection tool that maps PacBio HiFi reads back to the assembly and flags collapsed, misassembled, or low-confidence regions from coverage and secondary-allele signal. After NucFlag filtering, 4,650 centromeres were retained for downstream analysis (Supplementary Figure S10A, B). The acrocentric chromosomes and chromosome Y had too few haplotypes remaining after filtering to support robust per-chromosome clustering (chr13 *n* = 17, chr14 *n* = 12, chr15 *n* = 8, chr21 *n* = 14, chr22 *n* = 33, and chrY *n* = 8; Supplementary Figure S10B) and were excluded from this analysis. The remaining 18 non-acrocentric autosomes and chromosome X each retained sufficient haplotypes for analysis. For each of these chromosomes, haplotypes were grouped by their KaryoScope-derived satellite composition profiles using hierarchical clustering. Within each chromosome, the cluster containing the most haplotypes was designated major and all remaining clusters, including singletons, were designated minor. A single representative was selected per cluster, and these representatives were assembled into a unified pangenome dendrogram (Figure 5A; see Methods).

**Fig. 5.**
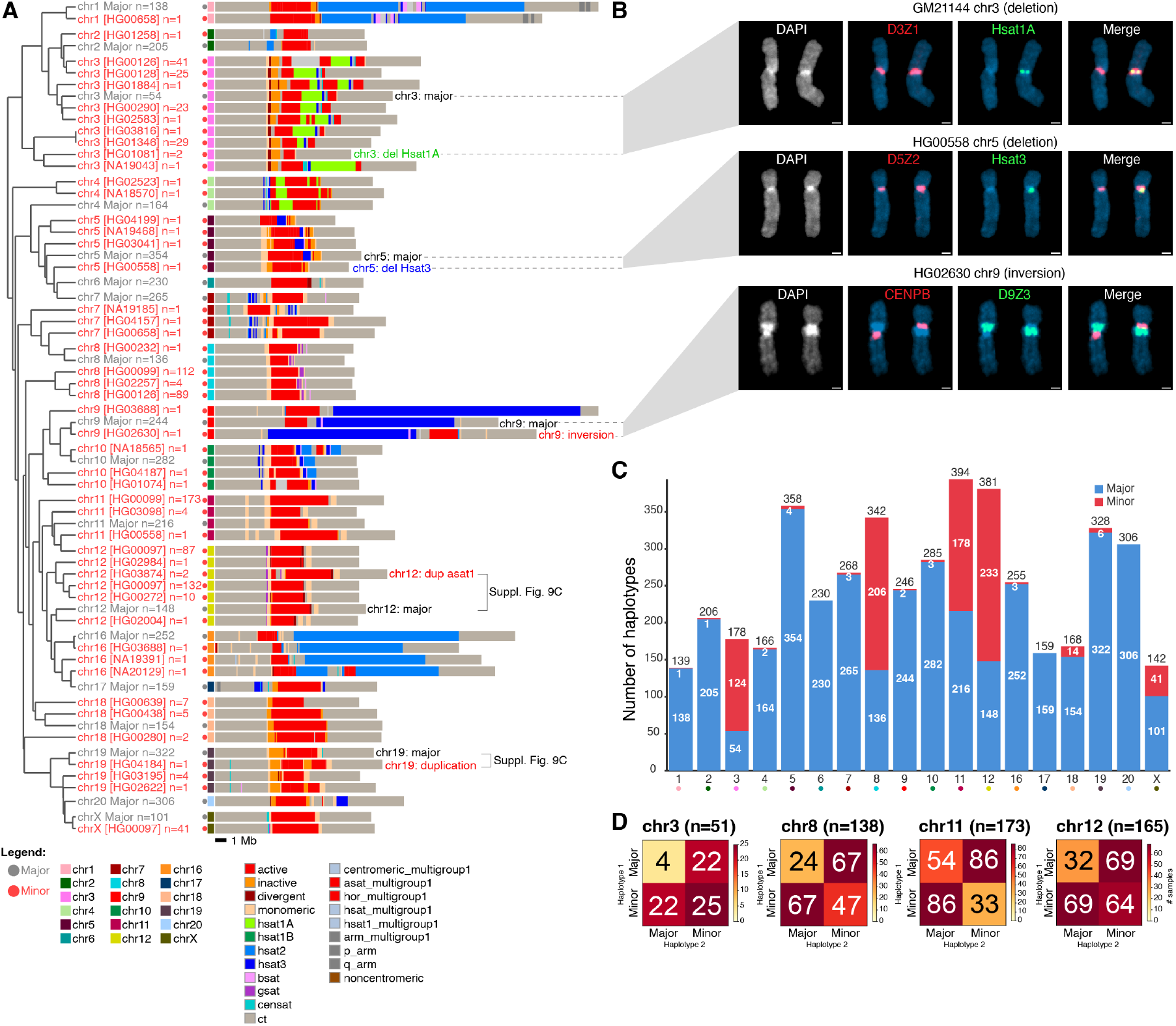
A pangenome-scale census reveals extensive centromeric satellite polymorphism, including ultra-rare variants. **(A)** Pangenome dendrogram of HPRC centromere cluster representatives. For each chromosome, haplotypes were clustered by their KaryoScope-derived satellite composition profiles and a single representative was selected per cluster (major, gray dot; minor, red dot; see Methods). Each row displays the KaryoScope annotation of one representative, coloured by satellite feature (legend, bottom left). Each row is labelled with chromosome, HPRC sample ID, and cluster size *n* (number of haplotypes sharing this compositional profile). Text callouts highlight the variants subsequently validated by FISH (panel B; Supplementary Figure S10C). **(B)** Fluorescence *in situ* hybridization (FISH) validation of minor compositional subtypes identified in panel A. Top: GM21144 (GIH) chr3 probed with D3Z1 (red, centromeric *α*-satellite) and HSat1A (green). Middle: HG00558 (CHS) chr5 probed with D5Z2 (red, centromeric) and HSat3 (green). Bottom: HG02630 (GWD) chr9 probed with CENPB (red) and D9Z3 (green). DAPI counterstain, individual channels, and merged images are shown. **(C)** Stacked bar chart of the number of HPRC haplotypes assigned to the major (blue) and minor (red) groups for each chromosome. **(D)** Diploid contingency tables of major/minor haplotype pairings for the four chromosomes with the largest minor-class representation: chr3 (*n* = 51 individuals), chr8 (*n* = 138 individuals), chr11 (*n* = 173 individuals), and chr12 (*n* = 165 individuals). Each *n* refers to individuals retaining both centromeres after quality filtering.

The dendrogram revealed a highly structured landscape. Centromere composition varied substantially across chromosomes, consistent with the known chromosome-specificity of *α*- satellite HOR families and the chromosome-biased distribution of HSat arrays. Within each chromosome, minor haplotypes departed from the major consensus through subtype-specific satellite gains, losses, or rearrangements. Some minor clusters contained only a single haplotype (*n* = 1), demonstrating sensitivity to structural variants present in as few as one of several hundred assemblies. This sensitivity is beyond the reach of consensus-matching annotation tools, which aggregate low-abundance signal into the dominant repeat call.

To distinguish genuine polymorphisms from potential assembly artifacts, we performed orthogonal FISH validation on three minor subtypes identified in the pangenome dendrogram (Figure 5B): an HSat1A deletion on chromosome 3 (GM21144), an HSat3 loss on chromosome 5 (HG00558), and a megabase-scale HSat3 inversion on chromosome 9 (HG02630). Each experiment used probes targeting the satellite subfamily implicated by KaryoScope rather than generic centromere markers, allowing direct visualization of the predicted compositional change at metaphase. In all three cases, the FISH signal pattern confirmed the KaryoScope-derived prediction. Two further minor subtypes, a chr12 D12Z3 variant (HG03816) and a chr19 D19Z3 variant (GM18960), showed comparable concordance (Supplementary Figure S10C). These five validated cell lines spanned five HPRC subpopulations (GIH, CHS, GWD, BEB, and JPT), representing South Asian, East Asian, and African continental ancestries, indicating that KaryScope’s detection sensitivity extends to haplotypes ancestrally distant from the T2T-CHM13v2.0 reference. To test whether these variants represented heritable germline polymorphisms rather than somatic or culture-acquired rearrangements, we extended the chromosome 9 validation to parental FISH of the HG02630 trio (Supplementary Figure S10C). The HSat3 inversion was present on the paternal homolog and absent from the maternal configuration, consistent with single-parent inheritance through meiosis. Together, these orthogonal validations established that KaryoScope’s compositional annotations capture genuine centromere structural variation at single-haplotype resolution, and that the HPRC Release 2 assemblies faithfully preserve this variation.

Although each minor subtype was individually rare, centromere composition was highly polymorphic in the aggregate. On chromosome 3, for example, nine distinct minor subtypes together accounted for 124 of 178 haplotypes (70%), with individual cluster sizes ranging from 1 to 41 (Figure 5A). Chromosomes 8, 11 and 12 similarly carried extensive minor populations (206 of 342, 178 of 394, and 233 of 381 haplotypes, respectively), distributed across multiple subtypes (Figure 5C). Other chromosomes carried narrower minor populations or none, with chr6, chr17, and chr20 represented entirely by major haplotypes.

We next asked whether this diversity was reflected within individuals. If polymorphism were partitioned between rather than within individuals, diploid genotypes would be dominated by major/major homozygotes. Instead, on the four chromosomes with the largest minor populations (chr3 *n* = 51, chr8 *n* = 138, chr11 *n* = 173, and chr12 *n* = 165 individuals with complete diploid genotypes), heterozygous major/minor pairings were the most common diploid configuration (Figure 5D). Compositional polymorphism therefore routinely pairs distinct centromere haplotypes on the two homologs of a single individual, suggesting that centromere heterozygosity is common rather than exceptional at these loci. Together, these observations place centromere satellite composition among the most structurally polymorphic regions of the human genome, and establish KaryoScope as a framework for cataloging that polymorphism at the haplotype and subfamily resolution required to connect it to function.

## Discussion

KaryoScope brings unified, base-pair resolution annotation to the entire genome, including the centromeres, subtelomeres, and acrocentric short arms that the pangenome era has made sequence-resolved but not yet interpretable at scale. Applied to the HPRC Release 2 pangenome, it resolves the sequence basis of Robertsonian translocations, the structural diversity of D4Z4 macrosatellites relevant to FSHD1, and the population-scale landscape of centromere polymorphism. These analyses establish the tool’s biological utility, but its architecture has implications well beyond satellite annotation.

KaryoScope extends a lineage of *k*-mer-based, alignment-free methods that have become foundational infrastructure in adjacent areas of genomics. Kallisto (26) and salmon (27) brought *k*-mer-based methods to transcript quantification; Kraken 2 (28) brought them to taxonomic classification. In each case, an input read is assigned to a single category. KaryoScope generalizes the paradigm: the same *k*-mer can carry labels across multiple feature sets simultaneously, so a single position in a sequence may be annotated as belonging to a specific chromosome, satellite family, repeat class, and gene at the same time. This shift, from sequence ( ∼3 Gb per haplotype) to a set of feature labels per position, renders the genome interpretable at scale, providing the kind of compact, labelled representation on which AI models can be trained.

The six feature sets released here illustrate the flexibility of the architecture. Any annotation that tiles a reference of interest can serve as a feature set, including FISH-probe libraries for *in silico* hybridization and breakpoint catalogs for recurrent-translocation detection. We are actively developing additional databases in this framework, including a chromosome-cytoband feature set, a GENCODE-based gene feature set, and a metagenomic database built from the Ref-Seq archaeal, viral, and bacterial collections, to be released in subsequent versions. Building *k*-mer databases from the human pangenome itself, rather than from T2T-CHM13v2.0 alone, will further allow KaryoScope to identify haplotype-, individual-, and parent-of-origin–specific *k*-mers and to track these signals across populations.

KaryoScope is well-suited to the structural complexity of cancer genomes. Applied to the HG008 pancreatic tumour assembly (10), KaryoScope resolved reciprocal translocations, dicentric chromosomes, chromoplexy, and breakpoints localized to the *α*-satellite region, features of complex variation in cancer that are difficult to interpret from variant calls alone. The same query architecture extends naturally to read-level data, where each read can be classified and quantified across the same feature sets used for assemblies. Because *k*-mer lookup is alignment-free, this is particularly valuable for reads spanning satellite arrays, structural rearrangements, or other repetitive regions where alignment is unreliable, and especially for clinical sequencing that typically yields reads rather than assemblies. KaryoScope is especially complementary to variant callers such as Severus (29): where those tools detect that a variant exists, KaryoScope can annotate the reads supporting that variant to help reveal the mechanism that produced it. A detailed read-level application to complex telomeric structural variants will appear in a forthcoming manuscript.

KaryoScope is built on the curated annotations it inherits: its *k*-mer database is derived from the RepeatMasker (12) and CenSat (15) annotations of T2T-CHM13v2.0, which it projects to assembly scale rather than supplanting. Several alignment-free frameworks for repeat and centromere analysis have emerged in parallel, each operating on a different signal: AniAnn’s (30) identifies tandem-array boundaries *de novo* from pairwise nucleotide identity, Centeny (17) uses CENP-box spacing to construct chromosome-specific centromere maps, and Shiraishi and colleagues (31) cluster *α*-satellite HOR haplogroups from rare *k*-mers. KaryoScope is distinguished in this landscape by providing a unified, base-pair resolution annotation across satellite families, interspersed repeats, and chromosome of origin in a single pass.

KaryoScope has limitations that frame the scope of its current applications. First, its annotations are projected from a reference-derived database; smoothing recovers most divergent *k*-mers when applied to assemblies from distant populations, but performance on samples with substantially novel satellite content will improve as pangenome-derived databases supplant single-reference ones. Second, KaryoScope’s resolution tracks assembly quality, a dependence we exploit by flagging contigs whose *k*-mer composition deviates strongly from reference expectations and letting the tool double as a quality-control instrument for assemblies. The current limits of this dependence are visible in our centromere analysis, where several acrocentric autosomes and chromosome Y were excluded because too few HPRC haplotypes survived quality filtering, reflecting the unfinished state of these regions. Third, the current database is built at a single *k*-mer length; applications targeting larger satellite repeat units may benefit from the variable-length indexing developed in our parallel HKS index (20).

As pangenome assemblies multiply across species and across human populations, alignment-free annotation tools will be essential for connecting sequence composition to function in the genome’s most variable regions. We offer KaryoScope as a foundation for that effort, and provide a pre-built database for the human genome alongside the tool.

## Methods

### Construction of KaryoScope feature sets

#### Overview

KaryoScope organizes genomic information through feature sets. A feature set is a complete labelling of the *k*-mers in the database, in which every *k*-mer is assigned to exactly one feature according to a given crierion (e.g., chromosome of origin, repeat family, or satellite identity). A feature set is constructed in two stages. First, every position in the reference genome (T2T-CHM13v2.0) is assigned to exactly one label, for example, chr1 in a chromosome-based feature set, LINE or SINE in a repeat-based feature set, or active_hor or HSat2 in a centromeric satellite feature set, producing a partition of the genome into non-overlapping, contiguous regions that tile each chromosome end to end. Second, each *k*-mer of the reference is assigned to exactly one feature. *K*-mers occurring at genomic positions corresponding to only a single label are assigned to a label-specific feature for that label; *k*-mers occurring at positions corresponding to multiple labels are assigned a multilabel feature corresponding to the least common ancestor (LCA) of those labels in a user-provided label hierarchy. The label hierarchy is a tree whose leaves are the labels and whose internal nodes group related labels (e.g., by biology or genome organization).

#### Gap filling

The annotation sources underlying a feature set typically cover only a fraction of the genome; for instance, the RepeatMasker annotations cover only repeat-containing positions, leaving non-repetitive sequence unannotated. Thus, producing a complete partition of the genome requires gap filling: every unannotated position must be assigned a label that reflects its genomic context. This gap filling procedure is the same for all the feature sets, differing only in the label used. For example, in the repeat feature set, all annotation gaps receive a single “nonrepeat” label; in the region feature set, gaps are labelled based on their position relative to the centromere (see below). After gap filling, adjacent regions sharing the same label are merged to produce the final partition of the genome.

#### Six feature sets

We constructed six feature sets from annotations of the T2T-CHM13v2.0 assembly (excluding chrM): a repeat feature set which labels the genome by repeat family, a region feature set which labels the genome by centromeric satellites, a chromosome feature set which labels the genome by chromosome, a subtelomeric feature set which labels features typically found on the subtelomeres, a gene feature set which labels the boundaries of genes, and an acrocentric feature set which labels features typically found on the p arms of the acrocentric chromosomes.

#### Repeat feature set

Repeat annotations were obtained from the T2T-CHM13v2.0 RepeatMasker v4.1.2p1 track hosted at the UCSC Genome Browser (https://hgdownload.soe.ucsc.edu/gbdb/hs1/t2tRepeatMasker/chm13v2.0_rmsk.bb), converted from bigBed to BED format with `bigBedToBed` (32). Each RepeatMasker entry encodes a repeat in the format name#class/subfamily; we extracted the class field as the label used by KaryoScope. The ambiguous “DNA?” class was merged into DNA, and the remaining fifteen class labels, DNA, LINE, LTR, SINE, RC, Retroposon, Satellite, rRNA, scRNA, snRNA, tRNA, srpRNA, Simple_repeat, Low_complexity, and Unknown, were retained at the class level.

The BED file containing these labels was sorted into genome order using samtools (33). Because RepeatMasker can produce overlapping annotations, where the same genomic position is annotated as multiple distinct repeat classes, each base of the genome was assigned a single label using a single-pass, priority-based flattening procedure. The priority order (highest to lowest) is rRNA, scRNA, snRNA, srpRNA, tRNA, Satellite, RC, Retroposon, DNA, LTR, SINE, LINE, Simple_repeat, Low_complexity, Unknown. This ordering favours rare, informative labels over abundant, low-information ones, so that the small number of bases occupied by, for instance, a tRNA repeat are not absorbed by an overlapping LINE annotation. Adjacent regions sharing the same flattened label were merged, and the gaps between annotated repeats were filled with the “nonrepeat” label, yielding a complete tiling of the genome.

#### Region feature set

Centromeric satellite annotations were obtained from CenSat v2.1 (https://raw.githubusercontent.com/hloucks/CenSatData/refs/heads/main/CHM13/chm13v2.0.cenSatv2.1.bed). Each CenSat entry contains one of the following 14 labels used by KaryoScope: active_hor, dhor, hor, mixedAlpha, mon, bSat, cenSat, ct, gSat, HSat1A, HSat1B, HSat2, HSat3, rDNA. Adjacent regions sharing the same label were merged. Because the underlying CenSat annotations are themselves non-overlapping, no priority-based flattening was required. Gap filling in this feature set is context-dependent rather than uniform: any unannotated region occuring before the centromere was filled with the “p_arm” label, any unannotated region occuring within the centromere was filled with “noncentromeric” (to indicate that it is not annotated in the centromeric feature set), and any unannotated region occuring after the centromere was filled with the “q_arm” label.

#### Chromosome feature set

Chromosome labels correspond directly to the T2T-CHM13v2.0 sequence names. Each position of the reference belongs to exactly one chromosome, so the chromosome labels are inherently non-overlapping and tile the genome end to end without requiring priority-based flattening or gap filling.

#### Subtelomeric feature set

This feature set consists of four labels, canonical telomere, noncanonical telomere, TAR1, and interstitial telomeric sequence (ITS), derived from a combination of de novo telomere detection and the T2T-CHM13v2.0 RepeatMasker v4.1.2p1 annotation.

Telomeric regions of the T2T-CHM13v2.0 assembly were first identified using `seqtk telo` (https://github.com/lh3/seqtk), which detects terminal arrays of the canonical (TTAGGG)n hexamer at chromosome ends. The FASTA sequence of each telomeric region was extracted with `bedtools getfasta` (34). Each position *p* within a telomeric region was assigned to the canonical_telomere label if the 31-mer beginning at *p* exactly matched one of the twelve canonical telomere 31-mers, the six in-frame rotations of (TTAGGG)n and the six rotations of its reverse complement (CCCTAA)n, and to the noncanonical_telomere label otherwise. Adjacent positions sharing the same label were merged.

TAR1, a subtelomeric satellite repeat, was identified by selecting all entries from an earlier release of the T2T-CHM13v2.0 RepeatMasker annotation (https://s3-us-west-2.amazonaws.com/human-pangenomics/T2T/CHM13/assemblies/annotation/chm13v2.0_RepeatMasker_4.1.2p1.2022Apr14.bed) whose repeat name contained the substring TAR1. The selected entries were sorted into genome order using samtools, and TAR1 hits within 1 kb of one another were merged into single intervals and labelled TAR1. The 1 kb threshold was chosen to consolidate the fragmented per-element hits that RepeatMasker reports across each contiguous TAR1 array into a single feature.

Interstial telomeric sequences (ITSs), internal copies of the (TTAGGG)n hexamer not contiguous with the chromosome termini, were identified in three steps. First, all entries from the same RepeatMasker release with class Simple_repeat whose repeat name contained any of the hexamer rotations of TTAGGG or CCCTAA were extracted. Second, the resulting candidate set was filtered to remove entries overlapping a `seqtk telo`-defined telomeric region, retaining only TTAGGG arrays interior to the chromosome. Third, surviving hits within 1 kb of one another were merged into single intervals and labelled ITS, using the same 1 kb threshold and rationale as for TAR1.

The four resulting BED files (canonical_telomere, noncanonical_telomere, TAR1, ITS) were concatenated and sorted into genome order. Because TAR1 elements frequently extend from the subtelomere into the telomeric array itself, the TAR1, canonical_telomere, and noncanonical_telomere annotations can overlap. Overlaps were resolved using the same single-pass, priority-flattening procedure used for the repeat feature set, with priority order (highest to lowest): canonical_telomere, noncanonical_telomere, TAR1, ITS. This ordering ensures that positions within true telomeric arrays, and within TAR1 elements that extend into them, are classified preferentially as telomere. Adjacent regions sharing the same flattened label were merged, and the remaining unannotated positions of each chromosome were filled with the nonsubtelomeric label.

#### Gene feature set

Gene annotations were obtained from the NCBI RefSeq annotation (35) of T2T-CHM13v2.0 (assembly accession GCF_009914755.1; https://ftp.ncbi.nlm.nih.gov/genomes/all/GCF/009/914/755/GCF_009914755.1_T2T-CHM13v2.0/GCF_009914755.1_T2T-CHM13v2.0_genomic.gtf.gz). The annotation is distributed in GTF format with sequences identified by NCBI accession (NC_060925.1 through NC_060948.1). These were renamed to the UCSC-style chromosome names (chr1-chr22, chrX, chrY).

Each position of the reference was assigned to exactly one of three labels, exon, intron, or intergenic, by considering all annotated transcripts simultaneously. Exon intervals were taken directly from exon records in the GTF, and intron intervals were derived for each transcript as the maximal intervals between consecutive exons of that transcript. Because a single position can be exonic in one transcript and intronic in another (for example, in alternatively spliced genes), we resolved such conflicts with a fixed precedence of exon > intron > intergenic: a position was labelled exon if it fell within any annotated exon of any transcript, intron if it fell within any annotated intron of any transcript but no exon, and intergenic otherwise. The classification was computed with a sweep-line procedure that maintains running counts of overlapping exon and intron intervals at every annotation boundary, producing a complete partition of each chromosome into non-overlapping exon, intron, and intergenic regions. GTF coordinates were converted from 1-based inclusive to the 0-based half-open BED convention. Adjacent regions sharing the same label were merged to produce the final BED file.

#### Acrocentric feature set

A BED file containing the locations of the distal junction (DJ) of the rDNA array, the proximal junction (PJ) of the rDNA array, the main rDNA array, the pseudohomologous regions (PHRs), and the SST1 macrosatellite in the T2T-CHM13v2.0 assembly was used as input to build the acrocentric feature set. These five labels collectively annotate the principal repeat and segmental-duplication features of the short arms of the five human acrocentric chromosomes (chr13-chr15, chr21, chr22).

Because the input annotations overlapped, each base of the genome was assigned a single label using the same singlepass, priority-based flattening procedure used elsewhere, with priority order (highest to lowest): SST1, PJ, PHR, DJ, rDNA. Adjacent regions sharing the same flattened label were merged, and the remaining annotated positions of each chromosome, including all the non-acrocentric chromosomes in their entirety, were filled with the nonacrocentric label.

### Feature set hierarchies and smoothing algorithm

#### Label hierarchies

Each feature set is accompanied by a label hierarchy: a rooted tree whose leaves are the labels of the feature set and whose internal nodes group related labels. The hierarchy serves two purposes. First, it determines the feature assignments at build time of *k*-mers occurring at genomic positions corresponding to multiple labels: such a *k*-mer is assigned to the least common ancestor (LCA) of those labels in the hierarchy, producing a multilabel feature. *K*-mers occuring at positions corresponding to a single label are assigned a label-specific feature corresponding to a leaf in the hierarchy. Second, the hierarchy is consulted by the smoothing algorithm described below, which recovers specific labels in regions where multilabel or novel *k*-mers have produced low-specificity intermediate calls.

#### Smoothing algorithm

When KaryoScope annotates a query sequence by querying its constituent *k*-mers against the database, contiguous stretches of label-specific calls are often interrupted by short runs of multilabel *k*-mers or by novel *k*-mers absent from the database. For example, SNPs, small indels, or repeat-element variation between the query and the T2T-CHM13v2.0 reference introduce small runs of novel *k*- mers. To recover specific feature assignments across these interruptions, we apply a sequence-context-aware smoothing procedure that uses the label hierarchy and the labels of neighbouring *k*-mers. The algorithm identifies windows of consecutive intervals that follow a specific to general to specific trajectory in the hierarchy, corresponding to regions where specificity temporarily decreases and then recovers, and reassigns the interior intervals to the LCA of the window endpoints.

Window detection proceeds via a two-phase scan over the merged-interval representation of the query. Starting from a left anchor interval *i* _*j*_, the ancestor phase scans rightward and accepts each subsequent interval whose feature is an ancestor of the previously accepted interval’s feature in the hierarchy, while skipping unrelated intervals (those on a different branch of the hierarchy). The phase terminates when (i) an interval more specific than the current accepted feature is encountered, (ii) the gap between consecutive accepted intervals exceeds ‘max_gap’ (set to 1000 bp), or (iii) a skipped unrelated interval would induce an overlap with a subsequent window. The descendant phase then continues rightward from the last interval accepted by the ancestor phase (the “peak”), accepting each subsequent interval whose feature is a descendant of the previously accepted feature, under symmetric termination conditions. The final accepted interval *i*_*r*_ becomes the right anchor.

For each detected window [*i* _*j*_, …, *i*_*r*_], the algorithm computes *l* = *LCA*( *f* (*i* _*j*_), *f* (*i*_*r*_)) in the hierarchy and reassigns every interior interval *i*_*a*_( *j < a < r*) whose feature is a descendant of *l* to the feature *l*. Interior intervals whose features are not descendants of *l* (i.e., intervals on a different branch that were skipped over during scanning) are left unchanged. Because the reassignment rule depends only on the endpoint features and the structure of the hierarchy, the produce is symmetric: applying the algorithm to a sequence and to its reverse complement produces equivalent labellings.

Novel *k*-mers, those absent from the database, initially labelled with the root feature of the hierarchy, are treated uniformly with multilabel *k*-mers during smoothing: if flanking context supports a more specific assignment, a novel *k*-mer is reassigned to the supporting LCA. Finally, the ‘max_gap’ value of 1000 bp was chosen by evaluating the concordance between the smoothed chromosome feature annotations of the HG002 assembly and the corresponding reference-derived ground truth.

### Genome-wide assembly visualization

Genome-wide karyotype visualizations were generated using custom Python scripts built on the `drawsvg` library (https://github.com/cduck/drawsvg). For each chromosome, the two haplotypes are displayed side by side, mirror-imaged about the center. Reading outward from the mirror axis, each haplotype comprises a centromere, full chromosome, and sub-telomere track. This design enables direct visual comparison of centromeric and subtelomeric composition between haplotypes.

All three tracks share a common binning procedure. The base-pair resolution smoothed annotation for the relevant feature set is divided into fixed-width bins; within each bin, each feature is scored by the number of bases it covers, and the bin is labelled with the highest-scoring feature, subject to two priority rules. First, when the feature set has a label hierarchy, leaf features (the most specific terminal labels) take precedence over internal-node features: if any leaf feature has non-zero overlap in a bin, only leaf features compete, and internal-node features are excluded from consideration. This prevents a bin containing a small amount of a specific label and a larger amount of its parent label from being labelled with the less informative parent. Second, the `novel` label, assigned to positions whose *k*-mers are absent from the database, including any *k*-mer overlapping an N, wins a bin only when it covers more than half of the bin; otherwise the highest-scoring non-novel feature is selected, even if it has less total overlap that novel. This rule ensures that bins containing fragmented annotated features against a sparse background of novel *k*-mers are reported by their informative labels, while contiguous unannotated stretches, such as assembly gaps within the acrocentric rDNA arrays, are correctly rendered as `novel`. After labelling, adjacent bins sharing the same feature assignment are merged into a single interval for visual compactness.

The full chromosome track bins the annotations at 1 Mb resolution, rendered at a scale of 1.4 pixels per megabase, spanning each chromosome from end to end.

The centromere track bins the annotations at 100 kb resolution, spanning only the centromere region. The centromere region is defined using the region feature set annotation and starts with the first 1 Mb bin with majority centromeric content and ends with the last 1 Mb bin with majority centromeric content. Panel height is fixed at 70% of the chromosome track height, with the largest centromere receiving the maximum panel height and all others scaled proportionally. This yields a uniform zoom factor across all chromosomes (4.9× for HG002 at default settings). Trapezoidal connectors link the centromere boundaries on the chromosome track to the edges of the centromere zoom panel.

The subtelomere track bins the annotations at 100 bp resolution, spanning the terminal 25 kb of each chromosome arm. Panel height is fixed at 21.4% of the chromosome track height per arm, yielding a zoom factor of ∼2,158 × relative to the chromosome track. When the p-arm and q-arm subtelomere panels would overlap on short chromosomes, the p-arm panel is offset outward to prevent collision.

For XY samples, chrX and chrY are displayed as a single visual unit with their centromeres placed next to each other. Multi-scaffold haplotypes (arising from fragmented assemblies) are handled by assigning synthetic haplotype names (h1a, h1b, etc.) to each scaffold.

The combined figure arranges chromosomes in a grid layout (3 rows of 8: chr1–8, chr9–16, chr17–22+X+Y) with feature set hierarchy trees displayed below. Hierarchy trees are rendered as vertical taxonomic trees showing the classification structure of each feature set, with horizontal bars scaled logarithmically by the number of distinct *k*-mers in the database assigned to each feature. Figures were rendered as SVG and rasterized to PNG at 300 dpi using `rsvg-convert`.

### KaryoScope annotation of query assemblies

Genome assemblies were annotated using KaryoScope (commit 7c976e04) with the KS_human_CHM13_v2 *k*-mer database, which encodes the chromosome, region (satellite), subtelomeric, acrocentric, repeat, and gene feature sets derived from the T2T-CHM13v2.0 reference. Annotations were binned at 100 bp, 100 kb, and 1 Mb resolutions for downstream visualization and summarization. Hierarchy-aware smoothing was applied for all analyses except the D4Z4 classification pipeline, which used unsmoothed KaryoScope output to preserve fine-grained structure within *β* -satellite arrays.

### HPRC Release 2 assemblies

Diploid haplotype-resolved assemblies were obtained from HPRC Release 2 (3). For the D4Z4 analysis (Figure 4), only contigs labelled “CM” (telomere-to-telomere) were retained, yielding *n* = 249 chromosome 4 haplotypes from 154 unique individuals and *n* = 333 chromosome 10 haplotypes from 208 unique individuals, corresponding to 224 unique individuals contributing at least one assembled chromosome 4 or 10 haplotype to the analysis. For the HG002 reference analysis (Figure 1), the hifiasm-trio diploid assembly of HG002 (NA24385) was used.

### Robertsonian translocation samples and breakpoint refinement

Three Coriell cell lines carrying Robertsonian translocations, GM03417 (45,XX,rob(14;21)(q10;q10)), GM03786 (45,XX,rob(13;14)(q10;q10)) and GM04890 (45,XX,rob(13;14)(q10;q10)), were analysed using the Verkko assemblies previously published (5). Cytogenetic abnormalities were described using the International System for Human Cytogenomic Nomenclature (ISCN) (36). We identified fusion scaffolds and assigned ISCN cytogenetic notation using custom scripts that apply ISCN conventions to KaryoScope chromosome annotations, flagging scaffolds carrying contiguous chromosome-specific *k*-mer signatures from two different acrocentric chromosomes. The chromosome-assignment transition interval, the genomic interval between the last 100 kb window dominated by one fusion partner’s chromosome-specific *k*-mers and the first 100 kb window dominated by the other’s, was derived from the chromosome feature set as part of this analysis (a 100 kb window was classified as dominated by a fusion partner when that chromosome’s *k*-mer content covered ≥25% of the window and accounted for ≥50% of the chromosome-specific content within it). For visualization, the SST1 satellite block at each fusion site, identified from the acrocentric feature set, was used as the central reference point of the multi-level zoom plots, while the chromosome-assignment transition interval was preserved as initially derived. Zoom plots at multiple scales (20 Mb to 200 kb) were generated with custom plotting scripts, each centered on the SST1 block midpoint at the fusion site.

For breakpoint refinement, we performed a symmetric cumulative *k*-mer analysis. For each fusion scaffold, the analysis window was defined as the chromosome-assignment transition interval extended by 1 Mb on each side. Within this window, we computed two cumulative fraction curves at 100 bp resolution from the chromosome feature set: a left-cumulative fraction for one fusion partner (the proportion of that partner’s chromosome-specific *k*-mers in the window lying at or before each evaluated position) and a right-cumulative fraction for the other (the proportion lying at or after each evaluated position). SST1-specific *k*-mer content was similarly computed from the acrocentric feature set. Each curve therefore ranges from 0 to 1: the left-cumulative curve rises from 0 at the left edge of the window to 1 at the right edge, while the right-cumulative curve falls from 1 at the left edge to 0 at the right edge. The combined score for each position in the window was defined as the sum of the values of the two cumulative curves at that position, reaching a theoretical maximum of 2 at a perfectly clean breakpoint, where one chromosome’s *k*-mers lie entirely to the left of the breakpoint and the other’s entirely to the right. The position along the scaffold corresponding to the maximum combined score was taken as the inferred breakpoint, and the 99% plateau width was defined as the genomic interval over which the combined score remains within 1% of its maximum. For visualization, the combined score curve was smoothed with a Gaussian kernel (sigma scaled to the downsampling ratio) and downsampled to 500 points.

### D4Z4 array annotation and classification

The terminal 250 kb of the q-arm of each chromosome 4 or chromosome 10 haplotype was defined as the subtelomeric analysis window. Two KaryoScope feature set annotations were used as input to the classification pipeline: the region feature set (using *β* -satellite and centromeric transition (ct) annotations) and the gene feature set (using exon, intron, and intergenic annotations). TAR1 annotations from the subtelomeric feature set were additionally used to define the telomeric boundary of each haplotype for array length calculations.

*β* -satellite fragments separated by ≤400 bp were merged into *β* -satellite repeat (BSR) clusters, consistent with the canonical D4Z4 architecture in which each ∼3.3 kb repeat unit contains a single ∼2.3 kb BSR element. Each BSR cluster was annotated by computing the fraction of its length covered by exon, intron, and intergenic features. Clusters that were ≥1,500 bp in length with ≥50% exon fraction were retained as candidate array elements (exon-dense clusters).

Arrays were initially defined as contiguous runs of exon-dense BSR clusters. Two exon-dense BSR clusters separated by more than 10 kb, or by any intervening non-exon-dense BSR cluster, were placed in separate arrays. Adjacent arrays were then re-merged if the intervening region (i) spanned ≤4 kb (a buffered cutoff above the canonical ∼3.3 kb D4Z4 repeat-unit length, so any larger gap is treated as a genuine inter-array boundary), with the intervening non-exon-dense BSR cluster(s) showing non-zero exon content, or (ii) was predominantly composed of novel sequence ( ≥70%), exon content ( ≥60%), or their combination ( ≥80%), provided no 3 kb sliding window within the gap contained ≥30% intron plus intergenic content. These merge thresholds were empirically chosen to bridge cases where binning artifacts or unclassified sequence fragment a single D4Z4 array into two apparent arrays, while the intron/intergenic boundary check preserves genuine array boundaries. Array boundaries were further extended by incorporating exon-dense fragments within 1,500 bp of either edge of the array; trailing extension additionally stopped at the first intron-rich BSR cluster (indicative of a terminal *β* -satellite). Within each array, pattern-based element merging produced consistently-sized elements ( ∼2.3 kb), and the final repeat count was the number of merged elements.

Two terminal features were evaluated distal to each array. A terminal *β* -satellite was defined as a BSR cluster of ≥2,500 bp with *<*35% exon content within 3 kb of the last exon-dense element, serving as a structural proxy for the A-type allele (4qA or 10qA). A segmental duplication-enriched centromeric transition (ct) terminal, marking the B-type allele (4qB or 10qB), was defined as a contiguous region of ≥1,000 bp with ≥50% centromeric transition (ct) density within 5 kb of the array end, where ct fragments separated by ≤500 bp were first merged. To confirm pLAM presence at the sequence level, BLAST was used to search for the pLAM polyadenylation signal sequence within the BSR regions of each classified array. Hits were classified as functional (ATTAAA) or non-functional (ATCAAA).

Arrays were classified as canonical A-type (C+; 4qA or 10qA) if a terminal *β* -satellite was detected and the array contained ≥1 element; canonical B-type (C ; 4qB or 10qB) if only a centromeric transition terminal was present and the array contained ≥3 elements; or degraded if no terminal was detected and the array contained ≥3 elements. Arrays failing all three criteria (fewer than 3 elements without a *β* -satellite terminal, or fewer than 3 elements with only a ct terminal) were classified as fragments and excluded from downstream analysis. Haplotypes in which the first array began within 1 kb of the left edge of the subtelomeric analysis window were flagged as potentially truncated. Per-haplotype summary statistics (array count, per-array classification, element count, median and mean element size, and total *β* -satellite content) were computed. Haplotypes with multiple or degraded arrays were flagged in the output. Chromosome 4 haplotypes with ≤10 elements in a canonical (C+) array, where a functional pLAM polyadenylation signal was confirmed by BLAST (see below), were considered consistent with the FSHD1 allelic configuration; sequence-level pLAM confirmation is required for clinical interpretation.

For orthogonal validation of KaryoScope repeat-unit counts, we compared against an independently derived D4Z4 HMM count table for chromosome 4 (see below). Of the 249 chromosome 4 haplotypes annotated by KaryoScope, 239 carried a single D4Z4 array (`num_arrays` = 1) and were thus eligible for one-to-one comparison; intersecting these by contig identifier with the HMM count table yielded 231 haplotypes with paired KaryoScope and HMM counts, on which the Pearson correlation reported in Supplementary Figure S8B was computed.

### HMM-based D4Z4 repeat-unit counting

The terminal *DUX4* 3.3 kb repeat from CHM13 chr4:193,538,193–193,541,504 was extracted, formatted as a FASTA alignment, and used to build a hidden Markov model with HMMER version 3.4 using `hmmbuild` (37). The repeats were then annotated with `nhmmer –cpu 12 --notextw --noali --tblout genome.out D4Z4rep.hmm genome.fa.` HMMER output files were converted to BED format using the gawk script available at https://github.com/enigene/hmmertblout2bed/blob/master/hmmertblout2bed.awk, invoked as `gawk -v th=0.7 -f hmmertblout2bed.awk genome.out> genome.bed.` Resulting annotations were filtered to retain only hits on chromosome 4, and hits larger than 2 kb were counted using `awk`.

The pLAM sequence was obtained from (38) and annotated in the assemblies using BLAST 2.16.0+ (39). BLAST databases were constructed from each assembly with `makeblastdb -in genome.fa -dbtype nucl -out genome`, and `blastn` searches were performed using `blastn -db genomeDB -outfmt 6 -query pLAM.fa -num_threads 8 -out genome.out`.

Searches were restricted to the terminal 250 kb of the q-arm of chromosome 4. Functional polyadenylation sequences (containing the ATTAAA allele) were identified as complete 100% matches to the query pLAM sequence. BLAST output files were converted to BED format with awk, and presence/absence of functional pLAM sequences was determined using `bedtools intersect`.

The pLAM query sequence was:

~~~
ACCTGCGCGCAGTGCGCACCCCGGCTGACGTGCAA
GGGAGCTCGCTGGCCTCTCTGTGCCCTTGTTCTTC
CGTGAAATTCTGGCTGAATGTCTCCCCCCACCTTC
CGACGCTGTCTAGGCAAACCTGGATTAGAGTTACA
TCTCCTGGATGATTAGTTCAGAGATATATTAAAAT
GCCCCCTCCCTGTGGATCCTATAGA
~~~

### Pangenome-scale centromere clustering and major/minor allele classification

#### Centromere interval definitions

Centromere intervals were defined per haplotype from the 100 kb and 1 Mb binned KaryoScope region BED files using a coarse-to-fine bounding-box procedure. A search window was first computed per chromosome by taking the minimum start and maximum stop coordinates of all 1 Mb bins assigned to a centromere satellite class and extending these by 1 Mb on each side (clipped to chromosome start = 0). The centromere interval was then defined as the minimum start and maximum stop coordinates of all 100 kb bins assigned to a centromeric satellite class falling entirely within that window. The 100 kb pre-smoothed region feature set annotations within these intervals constitute the upstream input to the analyses below.

#### Pangenome aggregation and assembly QC

HPRC Release 2 assemblies were first restricted to T2T contigs (GenBank CM-prefix accessions). For each retained contig, we examined NucFlag (16) QC flag counts (`Col, Dup, Err, NNN, COLLAPSE, COLLAPSE_OTHER, COLLAPSE_VAR, MISJOIN`). Only contigs with zero erroneous, collapsed, or collapsed-with-variation flags (`Err =` 0, `COLLAPSE =` 0, `COLLAPSE_VAR =` 0) were retained, yielding 4,650 chromosomes (Supplementary Figure S10A, B). For each retained chromosome, the KaryoScope region feature set annotations within its centromere interval defined that chromosome’s centromere annotation. These per-chromosome annotations were concatenated across all chromosomes and all assemblies into a single pangenome BED file. Acrocentric chromosomes (chr13, chr14, chr15, chr21, chr22) and chromosome Y were clustered but excluded from all downstream figures, because two few haplotypes survived QC to support robust per-chromosome clustering. The 18 non-acrocentric autosomes plus chromosome X constitute the denominator for the analyses below.

#### Per-chromosome structural clustering

Centromere structural diversity was modeled per chromosome as a directed transition graph over satellite-class labels (excluding novel features). For each haplotype, adjacent satellite-class labels along the centromere interval were emitted as directional edges *A* → *B*, and per-haplotype edge counts were stacked into a chromosome-specific feature matrix. Counts were transformed with log(count + 1) and standardized to per-column *z*-scores. Pairwise Euclidean distances were computed per chromosome and clustered with Ward linkage. The number of clusters *k* was selected automatically over *k*∈ [2, 10] by maximizing the silhouette score (sub-sampling cap = 2,000 haplotypes; random seed = 42). Within each chromosome, the largest cluster was designated the major allele and all remaining clusters minor alleles, including singletons; raw and max-normalized Euclidean distances to the major centroid were stored alongside the cluster label for each haplotype.

#### Two-stage refinement

The per-chromosome cluster labels were refined in two stages before plotting (silhouette threshold = 0.5; centroid distance threshold = 5 standard deviations). *Stage 1 (silhouette gating):* the silhouette score at *k* = 2 was recomputed on each chromosome’s edge feature matrix; chromosomes scoring *<* 0.5 were collapsed back to a single major cluster, suppressing forced splits on chromosomes without genuine substructure. *Stage 2 (centroid distance threshold):* within each chromosome, the Euclidean distance of every major-cluster haplotype to the major centroid was computed, and any haplotype with distance exceeding mean + 5 SD was reclassified as a minor allele, recovering rare individual outliers missed by the silhouette-driven *k*-selection.

#### Pangenome dendrogram, major/minor barplot, and diploid co-occurrence

For the pangenome dendrogram (Figure 5A), one representative haplotype per refined cluster was selected by Euclidean distance to the major centroid: the haplotype with minimum distance for the major cluster, and the haplotype with maximum distance for each minor cluster. These 66 representatives across the 18 included chromosomes (one major plus zero or more minor clusters per chromosome) were re-embedded in a single global edge feature matrix using the same directional edge mode and log(count + 1)*/z*-score normalization, and clustered with Ward linkage on Euclidean distances. Each row was annotated with a satellite-composition bar drawn from the haplotype’s 100 kb pre-smoothed region BED using the standard satellite colour palette, and a vertical chromosome-identity strip indicating which chromosome each representative came from. Total major and minor haplotype counts per chromosome were rendered as a stacked bar chart (Figure 5C). For the diploid allele co-occurrence analysis (Figure 5D), haplotypes were paired per individual by parsing the haplotype identifier `sample#{1 2}#contig`; only individuals for which both haplotypes survived NucFlag QC were retained. For each of the four chromosomes with the largest minor populations (chr3, chr8, chr11, chr12), a co-occurrence matrix of refined cluster assignments was constructed for the resulting paired-haplotype sample set, distinguishing major/minor and minor/minor diploid configurations.

### Chromosome spreads and fluorescence *in situ* hybridization

Human lymphoblastoid cell lines GM21144, HG00558, HG02628, HG02629, HG02630, HG03816, and GM18960 were obtained from Coriell and cultured in RPMI 1640 (Gibco) with L-glutamine supplemented with 15% fetal bovine serum (FBS) at 37°C with 5% CO_2_. For preparation of chromosome spreads, cells were blocked in mitosis by addition of Karyomax colcemid solution (0.1 *μ*g/mL, Life Technologies) for 6–7 h and collected by centrifugation at ∼250*g* for 5 min. Collected cells were incubated in hypotonic 0.4% KCl solution for 12 min and prefixed by addition of methanol:acetic acid (3:1) fixative solution (1% total volume). Pre-fixed cells were spun down and then fixed in methanol:acetic acid (3:1).

For FISH, chromosome spreads were dropped on glass slides and incubated at 65°C overnight. Before hybridization, slides were dehydrated in a 70%, 80%, and 100% ethanol series for 2 min each. Slides were denatured in 70% deionized formamide / 2× SSC solution pre-heated to 72°C for 1.5 min. Denaturation was stopped by immersing slides in a 70%, 80%, and 100% ethanol series chilled to -20°C. Labeled DNA probes were denatured separately in hybridization buffer by heating to 80°C for 7 min before applying to denatured slides.

Fluorescently labelled human CenSat probes for D3Z1 (Cytocell, cat. #LPE003R), D1Z7/D5Z2/D19Z3 (cat. #LPE005R), D9Z3 (cat. #LPE019G), and D12Z3 (cat. #LPE012R) were obtained from Cytocell. The HSat1A_1 probe was generated by cloning a consensus monomer into the pUC-19 vector; it was labelled with biotin-16-dUTP using a nick translation kit (Enzo Life Sciences) and detected with streptavidin conjugated to Cy5 (Thermo Fisher). The HSat3 oligonucleotide consensus probe, labelled with FAM, was synthesized by IDT (5^′^ /56-FAM/(AATGG)_8_AAT/36-FAM/ 3^′^). The PNA CENPB probe labelled with Cy3 was obtained from PNA Bio (cat. #F3002).

Specimens were hybridized to probes under a HybriSlip hybridization cover (Grace Bio-Labs) sealed with Cytobond (SciGene) in a humidified chamber at 37°C for 48–72 h. After hybridization, slides were washed in 50% formamide / 2× SSC three times for 5 min per wash at 45°C, then in 1× SSC at 45°C for 5 min twice and at room temperature once. For biotin detection, slides were incubated with streptavidin conjugated to Cy5 (Thermo Fisher) for 2–3 h in PBS containing 0.1% Triton X-100 and 5% bovine serum albumin (BSA), then washed three times for 5 min with PBS / 0.1% Triton X-100. Slides were rinsed in water, air-dried, and mounted in Vectashield containing DAPI (Vector Laboratories).

Confocal Z-stack images were acquired on a Nikon Ti-E microscope equipped with a Plan Apo 100× oil-immersion objective (NA 1.45), a Yokogawa CSU-W1 spinning disk, an ORCA-Fusion BT sCMOS camera (Hamamatsu), and NIS-Elements software. All image processing was performed in ImageJ/Fiji. Maximum intensity projections of CenSat FISH images from spinning-disk confocal Z-stacks were generated. Chromosomes of interest were manually segmented, oriented vertically, and shown in composite mode.

## Software and code availability

KaryoScope is freely available as open-source software at https://github.com/barthel-lab/KaryoScope. HPRC Release 2 diploid assemblies analysed in this study are available at https://humanpangenome.org/hprc-data-release-2/. The three Robertsonian translocation assemblies for GM03417, GM03786, and GM04890 are available via dbgap from phs003920.v1.p1 in accordance with patient consent from Coriell.

## ACKNOWLEDGEMENTS

This work was supported by the Department of Defense (award HT9425-23-1-0844 to F.P.B.) and the Lennar Foundation. Large language models were used in the preparation of this manuscript to improve grammatical accuracy and readability, and to assist with the development of analysis code. The authors reviewed all edits and retain sole responsibility for the accuracy and integrity of the content.

## Supplementary Information

### Supplementary Note 1: Benchmarking details

#### T2T-CHM13v2.0 RepeatMasker self-benchmark

To validate the integrity of the KaryoScope repeat feature set, we annotated the T2T-CHM13v2.0 reference assembly and compared the output to RepeatMasker annotations. After smoothing, 98.69% of *k*-mer positions matched RepeatMasker exactly, with an additional 0.64% classified as compatible (Supplementary Figure S5A, C). Only 0.47% of positions were annotated with an incorrect repeat class. The vast majority of these involved the merged tandem-repeat category being classified as interspersed repeats, reflecting transposable-element fragments embedded within tandem-repeat regions: Repeat-Masker annotates such regions by their dominant repeat class, while KaryoScope identifies the TE-derived *k*-mers within them. An additional 0.21% of positions not annotated as repetitive by RepeatMasker were classified into a repeat family by KaryoScope, likely representing ancient or repeat-derived sequences below RepeatMasker’s detection threshold.

**Fig. S1.**
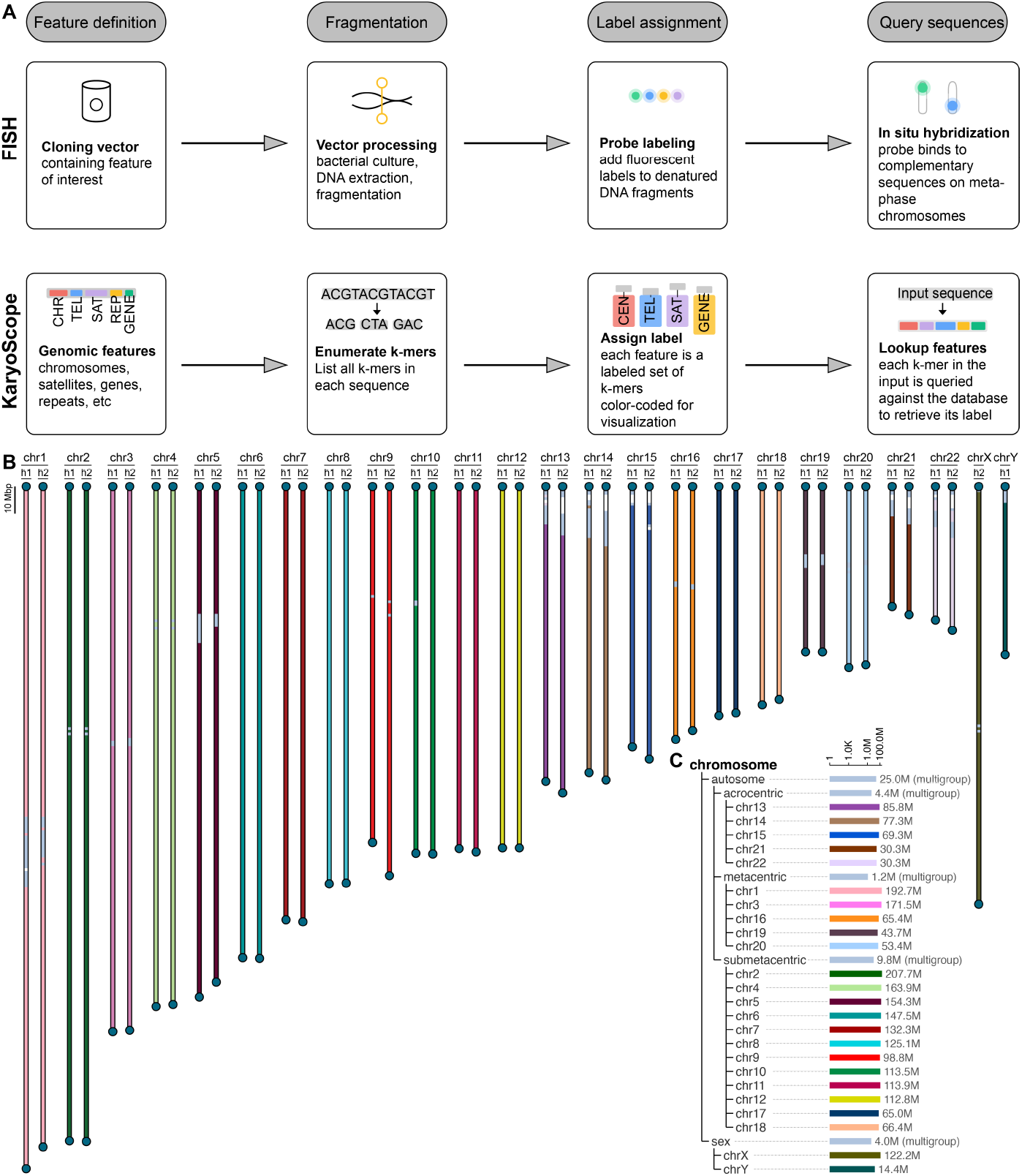
Conceptual comparison of FISH and KaryoScope karyotyping workflows. **(A)** Conventional CO-FISH workflow (top) requires cloning vectors, cell culture, DNA extraction, fragmentation, fluorescent labelling, and *in situ* hybridization to produce a cytogenetic karyotype, whereas KaryoScope (bottom) queries each *k*-mer in an assembled genome sequence against a labelled *k*-mer database to produce a computational karyotype. **(B)** Genome view of HG002 showing the chromosome feature set, with each chromosome coloured by chromosome of origin. All 24 chromosomes are displayed: both haplotypes (h1, h2) for the 22 autosomes, and a single copy each for chrX and chrY. **(C)** Chromosome feature set hierarchy tree showing log-scaled *k*-mer counts per chromosome, with autosomes grouped by morphology (acrocentric, metacentric, submetacentric) and shown alongside the sex chromosomes.

**Table S1.** Runtime and memory usage of KaryoScope-KMC annotation pipeline steps. Per-haplotype wall-clock (minutes) and peak memory (GiB, `max_rss`) for the two principal annotation steps, collected by the Snakemake `benchmark:` directive. For `smooth_features.py`, six jobs run in parallel per haplotype (one per feature set); “wall-clock (max)” reports the slowest feature set (critical path on a cluster), and “wall-clock (sum)” reports the total across feature sets (serial equivalent). All jobs used 16 threads on Intel Xeon Ice Lake 2.80 GHz nodes of the TGen Gemini cluster against the KS_human_CHM13_v2 database.

| Sample | Hap | Script | *n* jobs | Wall-clock<br>max (min) | Wall-clock<br>sum (min) | Peak RSS<br>(GiB) | CPU-time<br>(min) |
| --- | --- | --- | --- | --- | --- | --- | --- |
| T2T-CHM13v2.0 | hap1 | get_featureIDs | 1 | 4.58 | 4.58 | 33.96 | 35.88 |
| T2T-CHM13v2.0 | hap1 | smooth_features.py | 6 | 3.31 | 15.12 | 43.13 | 112.32 |
| HG002 | hap1 | get_featureIDs | 1 | 4.46 | 4.46 | 34.45 | 35.80 |
| HG002 | hap1 | smooth_features.py | 6 | 2.83 | 13.79 | 43.95 | 115.76 |
| HG002 | hap2 | get_featureIDs | 1 | 4.59 | 4.59 | 34.94 | 35.22 |
| HG002 | hap2 | smooth_features.py | 6 | 3.25 | 16.25 | 44.90 | 122.91 |

**Fig. S2.**
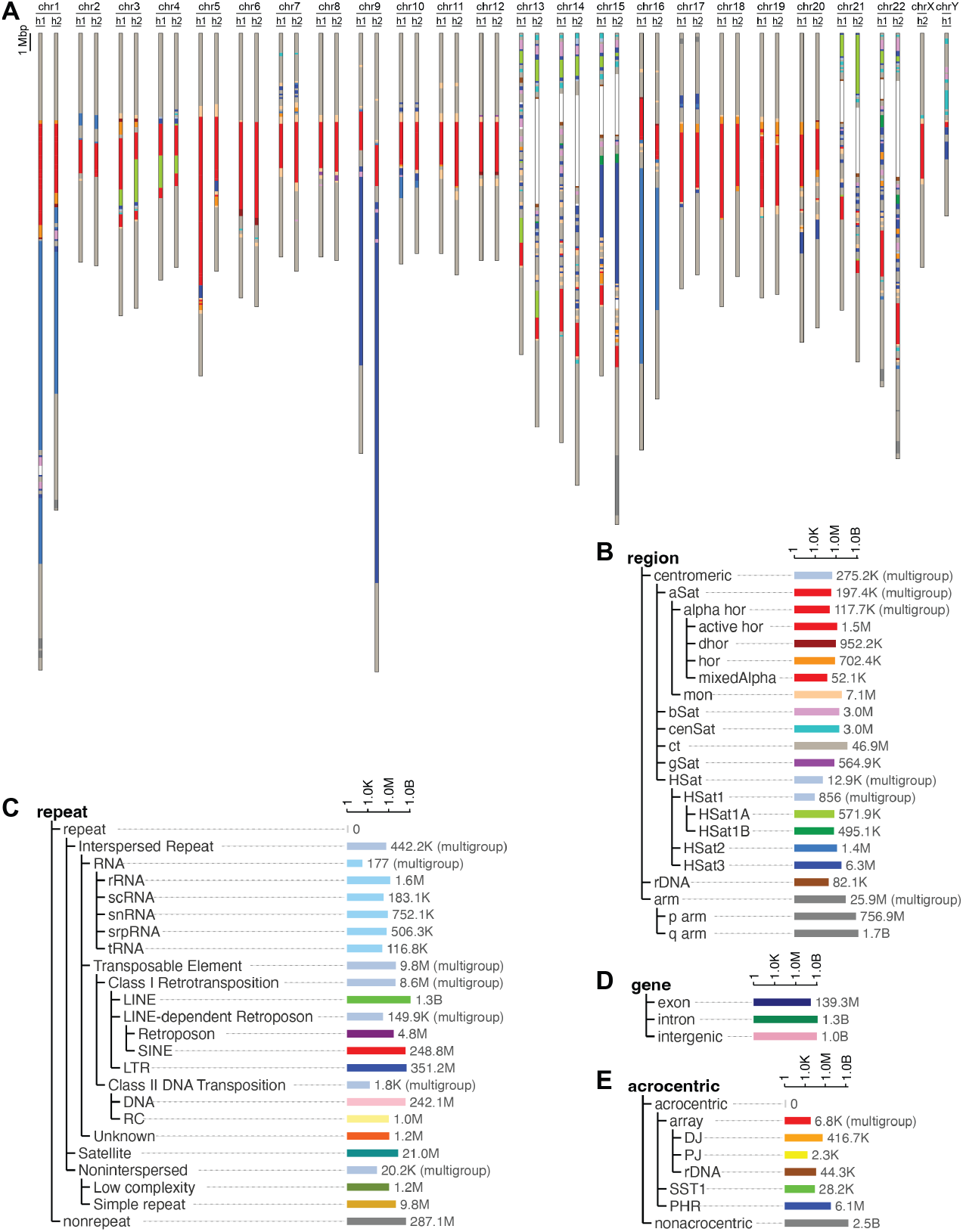
Centromere view of the HG002 diploid assembly. **(A)** Centromere view of HG002 showing the region (satellite) feature set for all chromosomes, highlighting haplotype- and chromosome-specific variation in satellite composition. Centromeric regions are visible as coloured segments showing satellite composition (HOR subfamilies including active, inactive, and divergent HORs; mixed and monomeric *α*-satellite; *β* -satellite; cenSat; centromeric transition (ct) regions; gSat; human satellite subfamilies including HSat1A, HSat1B, HSat2, HSat3; rDNA) against the gray chromosome arm background. **(B)** Region feature set hierarchy tree. **(C)** Repeat feature set hierarchy tree, showing major repeat classes including retrotransposons (LINE, SINE, LTR, Retroposon), DNA transposons (DNA, RC), tandem repeats (Satellite, Simple repeat, Low complexity), RNA (rRNA, scRNA, snRNA, srpRNA, tRNA), and unclassified elements. **(D)** Gene feature set hierarchy tree (exon, intron, intergenic). **(E)** Acrocentric feature set hierarchy tree, showing distal junction (DJ), proximal junction (PJ), rDNA, SST1 macrosatellite, and pseudohomologous region (PHR).

**Fig. S3.**
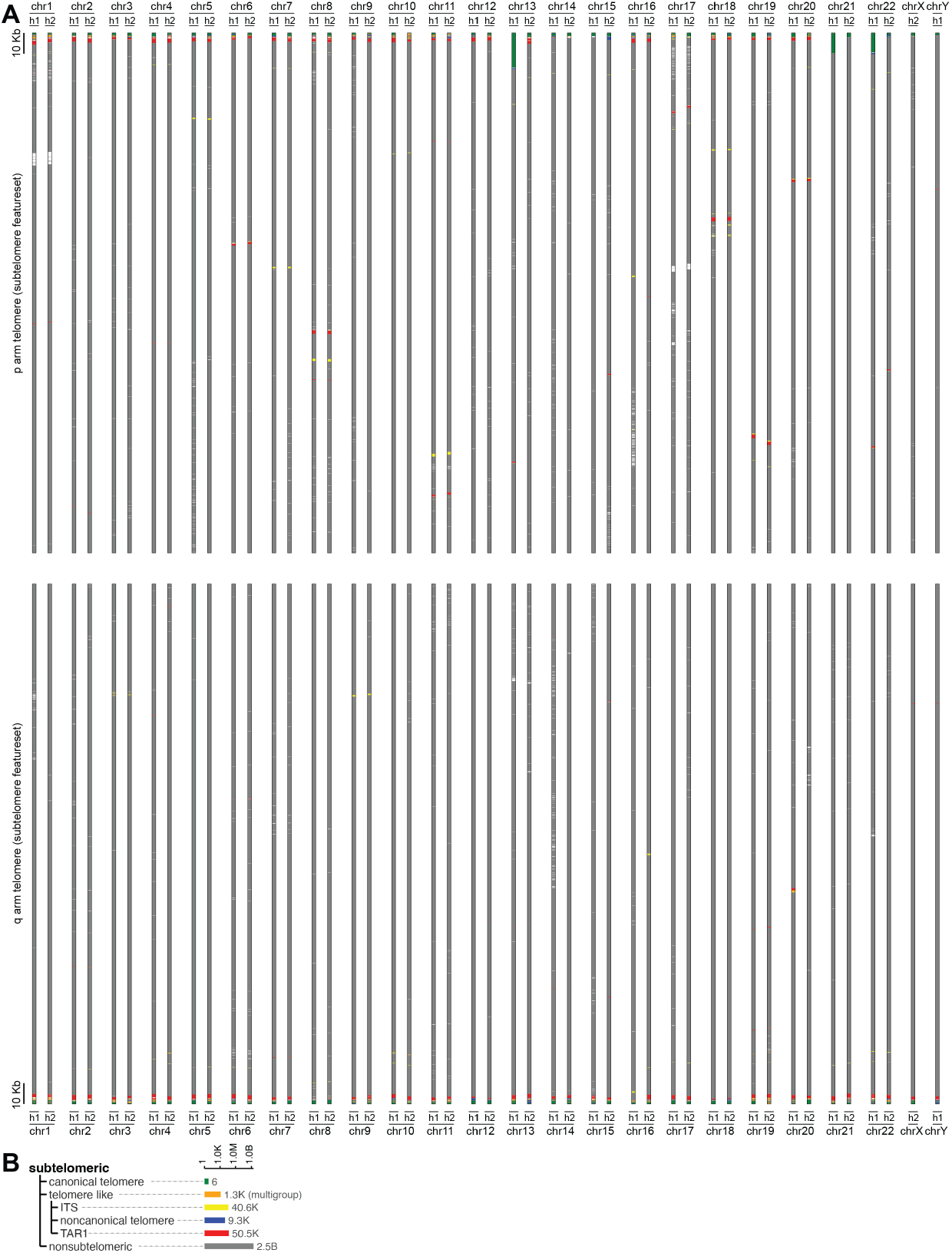
Subtelomeric view of the HG002 diploid assembly, subtelomeric feature set. **(A)** Terminal 250 kb of each chromosome arm coloured by the subtelomeric feature set. Both haplotypes (h1, h2) are shown for autosomes; chrX and chrY are shown singly. Top: p-arm telomeres. Bottom: q-arm telomeres. Canonical (TTAGGG)_*n*_ telomere tracts are present at all chromosome termini in both haplotypes. TAR1 elements are identified at most chromosome ends. Interstitial telomeric sequences and noncanonical telomere repeats are visible at select loci. Color coding for all subtelomeric features is given in panel B. **(B)** Subtelomeric feature set legend with *k*-mer counts.

**Fig. S4.**
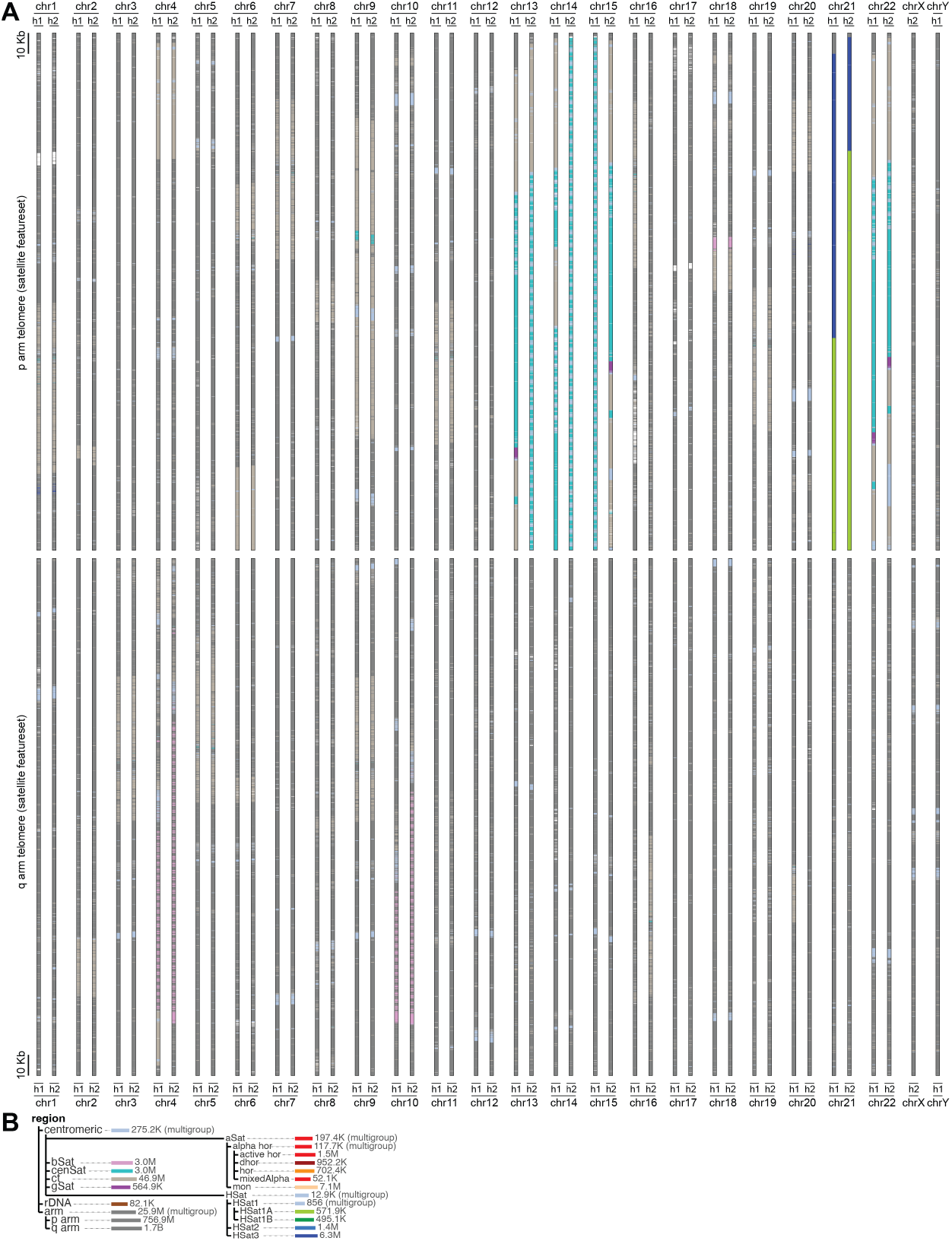
Subtelomeric view of the HG002 diploid assembly, region (satellite) feature set. **(A)** Same subtelomeric view as Supplementary Figure S3 but coloured by the region (satellite) feature set. Top: p-arm telomeres. Bottom: q-arm telomeres. Most subtelomeric regions are devoid of satellite content; the most prominent satellite signals are *β* -satellite arrays on the 4q and 10q subtelomeres (marking the distal boundary of the D4Z4 macrosatellite arrays) and acrocentric short-arm features (HSat, rDNA) at several p-arm termini. **(B)** Region feature set legend with *k*-mer counts.

**Fig. S5.**
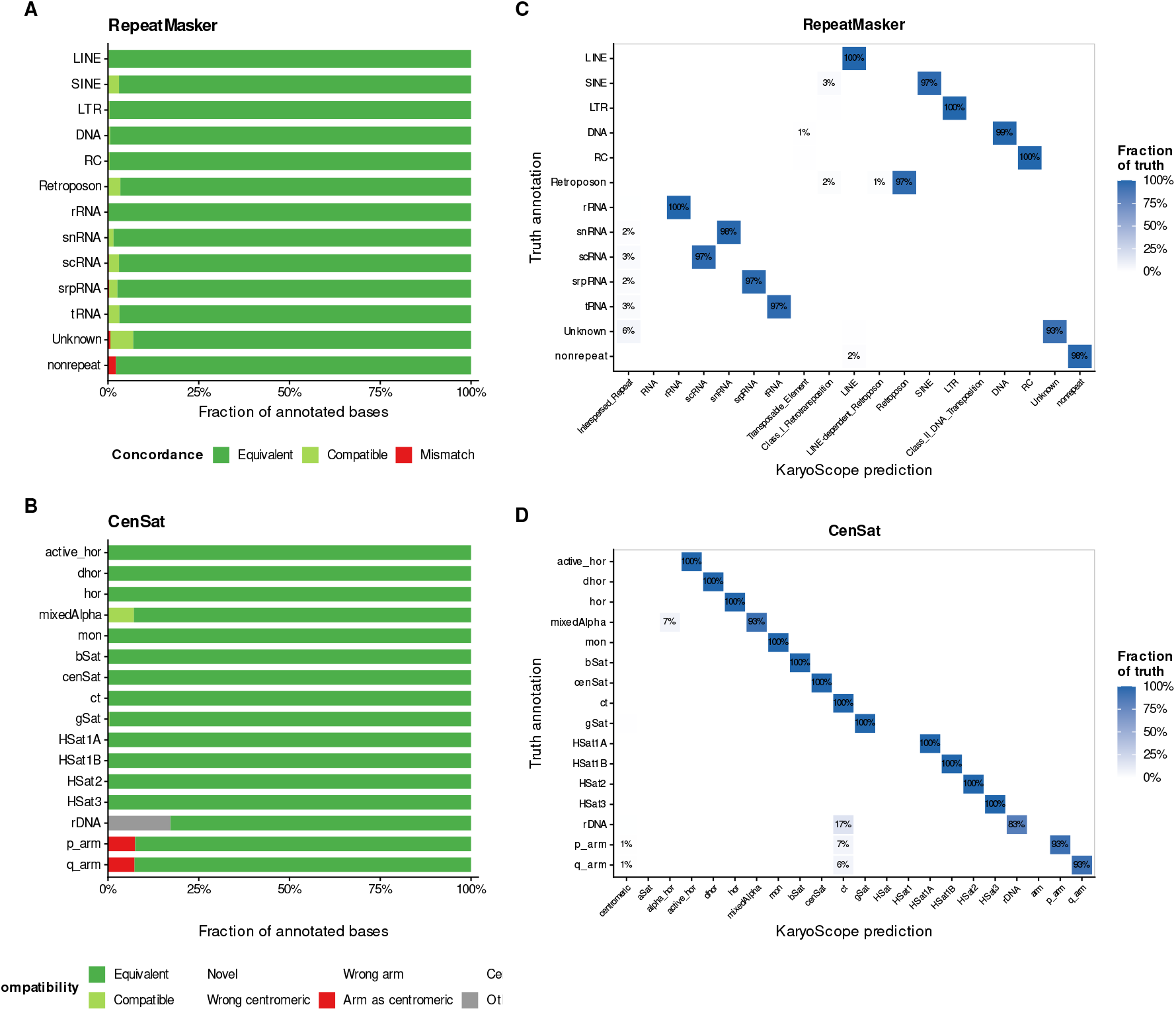
KaryoScope self-benchmark concordance on the T2T-CHM13v2.0 reference assembly. KaryoScope was applied to the T2T-CHM13v2.0 reference assembly (from which the database was constructed) and the resulting annotations were compared to the source RepeatMasker and CenSat v2.1 annotations as truth. The structure of this figure mirrors Figure 2, which shows the cross-genome benchmark on HG002. **(A)** Per-category concordance for the repeat feature set (RepeatMasker truth). Bars show the fraction of annotated bases per category that match the truth annotation exactly (Equivalent), at a less specific level in the feature hierarchy (Compatible), or as an incorrect class (Mismatch). **(B)** Per-category concordance for the region (centromeric satellite) feature set (CenSat truth), with mismatches further partitioned into wrong centromeric subtype, wrong arm, arm-as-centromeric, centromeric-as-arm, and other. **(C)** Confusion matrix showing KaryoScope predictions (columns) versus RepeatMasker truth (rows). Cell shading encodes the fraction of each truth category assigned to each predicted category. Brackets along the bottom axis indicate the hierarchical structure of the repeat feature set, allowing compatible parent-category predictions to be distinguished from outright mismatches. **(D)** Confusion matrix for the region feature set against CenSat truth, with hierarchical brackets for centromeric and arm subcategories.

Smoothing substantially improved classification specificity: prior to smoothing, 89.77% of *k*-mer positions exactly matched RepeatMasker and 9.56% were compatible, reflecting positions where *k*-mer evidence was insufficient to resolve the most specific repeat class. After smoothing, nearly all compatible assignments were resolved to their correct labels, increasing equivalent classifications to 98.69% while reducing compatible to 0.64%. Importantly, smoothing introduced only 264 kb (0.008%) of new misclassifications across all non-tandem-repeat categories, demonstrating that the smoothing algorithm overwhelmingly corrects rather than propagates errors. Within the merged tandem-repeat category, 93.87% of *k*-mer positions matched RepeatMasker exactly, with the remaining 6.13% annotated as interspersed repeat classes, predominantly LINE (5.1 Mb), SINE (3.8 Mb), and other Class I retrotransposons, consistent with the TE-derived *k*-mer effect described above.

#### T2T-CHM13v2.0 CenSat self-benchmark

To validate the KaryoScope region feature set, we annotated the T2T-CHM13v2.0 reference assembly and compared the output to CenSat v2.1 annotations. Within centromeric and rDNA regions (321.2 Mb), 99.44% of *k*-mer positions matched CenSat exactly after smoothing, with no incorrect centromeric subtype classifications: every satellite subtype, including active HOR, inactive HOR, monomeric *α*-satellite, and all human satellite families (HSat1A, HSat1B, HSat2, HSat3), matched at 99.8– 100% per-category accuracy (Supplementary Figure S5B, D). The only centromere-internal discordance was 1.7 Mb (0.53%) of rDNA annotated as centromeric, likely reflecting shared *k*-mer content between rDNA arrays on the acrocentric short arms and adjacent centromeric sequence. An additional 0.03% of centromeric positions were classified as compatible after smoothing.

Genome-wide, 93.33% of *k*-mer positions matched CenSat exactly after smoothing. Smoothing substantially reduced compatible classifications, from 5.66% to 0.005%, resolving nearly all less-specific annotations to their correct labels. The 6.61% discordance was almost entirely chromosome-arm positions annotated as centromeric (205.9 Mb), predominantly as ct (centromeric transition; 180.8 Mb, 88% of the total). The ct annotation in CenSat v2.1 marks non-satellite sequences interspersed between centromeric satellite arrays, largely composed of segmental duplications. These sequences share *k*-mer content with paralogous regions in the chromosome arms, causing KaryoScope to annotate arm positions with the ct label. Notably, the total amount of arm sequence annotated as centromeric was identical before and after smoothing (205.9 Mb), confirming that this is an inherent property of shared *k*-mer content rather than a smoothing artifact. Smoothing did, however, improve the specificity of these annotations, resolving 75.7 Mb of generic centromeric calls to the more informative ct label.

#### HG002 RepeatMasker cross-genome benchmark

To evaluate KaryoScope’s cross-genome repeat classification accuracy, we annotated the HG002 diploid assembly (haplotype 1, paternal; haplotype 2, maternal) using the T2T-CHM13v2.0-derived repeat database and compared the output to HG002’s own RepeatMasker annotations, processed through the same pipeline used to construct the database. Assembly gaps (stretches of Ns) were excluded from the evaluation, as *k*- mers overlapping these positions do not represent valid genomic sequence. This excluded 7.6 Mb from haplotype 1 and 23.4 Mb from haplotype 2, corresponding primarily to unassembled rDNA arrays on the acrocentric short arms, with larger arrays in the maternal haplotype. After exclusion, 2,940,260,101 *k*-mer positions were evaluated in haplotype 1 and 3,028,094,375 in haplotype 2.

After smoothing, 95.25% (haplotype 1) and 95.21% (haplotype 2) of *k*-mer positions matched RepeatMasker exactly, with an additional 1.06% and 1.08% classified as compatible, respectively (Figure 2A, C). These results were highly consistent between haplotypes, with per-category accuracy differing by less than 0.5 percentage points across all repeat classes. Smoothing substantially improved classification specificity: prior to smoothing, 84.6–84.8% of positions matched exactly and 9.2–9.5% were compatible; after smoothing, nearly all compatible annotations were resolved to their most specific labels while novel calls were reduced from 2.5–2.6% to under 0.05%. The major interspersed repeat classes each matched RepeatMasker at high rates, LINE (98.3%), LTR (97.7%), DNA (96.8%), and SINE (94.7%), with nearly identical values across both haplotypes. The modest reduction relative to the T2T-CHM13v2.0 self-benchmark (where these classes exceeded 99%) reflects genuine inter-individual variation in repeat content, including population-specific transposable-element insertions and sequence divergence at orthologous repeat loci.

Three categories showed notable cross-genome effects. First, 7.0% (haplotype 1) and 6.8% (haplotype 2) of positions lacking RepeatMasker annotations were annotated as a repeat family by KaryoScope, predominantly LINE and LTR. These likely represent repeat-derived sequences in HG002 that fall below RepeatMasker’s detection threshold but retain sufficient *k*-mer similarity to annotated repeats in T2T-CHM13v2.0. Second, 80.1% (haplotype 1) and 75.2% (haplotype 2) of positions in the merged tandem-repeat category matched RepeatMasker exactly, with 18.0–22.5% annotated as interspersed repeat classes, reflecting both the *k*-mer-sharing effects described in the T2T-CHM13v2.0 self-benchmark and well-documented inter-individual variation in satellite-array composition. Third, only 20.5% (haplotype 1) and 17.6% (haplotype 2) of positions in the Unknown repeat class matched RepeatMasker exactly, with 68.4–71.8% annotated as non-repetitive. Unknown repeats are by definition poorly characterized and highly diverged elements; the low cross-genome concordance indicates that HG002’s Unknown repeat *k*-mers are sufficiently divergent from T2T-CHM13v2.0’s to fall below the classification threshold. At 8.3 Mb total (0.3% of the genome), this category has negligible impact on overall accuracy.

#### HG002 CenSat cross-genome benchmark

To evaluate KaryoScope’s cross-genome centromeric satellite classification accuracy, we annotated the HG002 diploid assembly using the T2T-CHM13v2.0-derived region database and compared the output to HG002’s own CenSat v2.0 annotations. After smoothing, 92.84% (haplotype 1) and 92.73% (haplotype 2) of *k*-mer positions matched CenSat exactly genomewide (Figure 2B, D). As with the T2T-CHM13v2.0 self-benchmark, the dominant source of discordance was arm positions annotated as centromeric (6.6–6.8%), almost entirely reflecting the pericentromeric ct effect described above. The magnitude of this effect was virtually identical across T2T-CHM13v2.0 (6.6%) and both HG002 haplotypes (6.6–6.8%), confirming that it is a genome-wide property of shared *k*-mer content between ct regions and chromosome arms rather than a sample-specific artifact.

Restricting the analysis to centromeric and rDNA regions, 98.24% (haplotype 1) and 98.74% (haplotype 2) of *k*-mer positions matched CenSat exactly, with only a modest reduction from the T2T-CHM13v2.0 self-benchmark (99.44%). The majority of centromeric satellite subtypes each matched CenSat at high rates across both haplotypes: HSat1A (100%), HSat2 and HSat3 (99.9%), bSat (99.8%), cenSat (99.3–99.4%), active HOR (97.6–98.5%), and monomeric *α*-satellite (98.0–98.3%).

Only 0.015% of centromeric positions were annotated with an incorrect centromeric subtype, and these were concentrated within the *α*-satellite family, cross-annotations among active_hor, hor, dhor, mixedAlpha, and mon, where inter-individual sequence variation and inherent *k*-mer similarity among related subtypes make precise discrimination challenging. In particular, only 58% of positions in the mixedAlpha category (228–269 kb) matched CenSat exactly, with the remainder annotated as other *α*-satellite subtypes. The mixedAlpha category by definition represents hybrid *α*-satellite arrays containing elements of multiple subtypes, and this cross-annotation is expected given the composite nature of these arrays and does not represent a categorical error.

## Supplementary Note 2: Performance and scaling

We benchmarked the KaryoScope-KMC annotation pipeline on three haplotype assemblies (T2T-CHM13v2.0 hap1; HG002 hap1; HG002 hap2) against the KS_human_CHM13_v2 *k*-mer database on a cluster node (AMD EPYC 7302, 64-core, 3 GHz; 503 GiB RAM; 16 CPU threads per job). The two principal steps, *k*-mer-based feature assignment (`get_featureIDs`) and hierarchical feature smoothing (`smooth_features.py`), completed in mean wall-clocks of 4.54 ± 0.08 and 3.13 ± 0.26 minutes per haplotype respectively, with peak resident memory of 34.45 ± 0.49 GiB and 43.99 ± 0.89 GiB (*n* = 3; Supplementary Table S1). End-to-end per-haplotype wall-clock through both steps was 7.67 ± 0.33 minutes, demonstrating that a complete KaryoScope-KMC annotation of a human haplotype can be generated in under eight minutes on standard HPC hardware without specialized accelerators.

We additionally benchmarked two pipeline upgrades that substantially reduce runtime and memory beyond the released KaryoScope-KMC defaults (Supplementary Figure S6). For *k*-mer-based feature assignment, replacing the KMC back-end with the hierarchical *k*-mer set (HKS) index (20), a variable-length *k*-mer index built on the Spectral Burrows– Wheeler Transform (21), increased feature-lookup throughput on HG002 hap1 from 11.1 to 28.1 Mb/s at 16 threads (2.5 ×), while reducing peak resident memory from 33.4 to 10.9 GiB (3.1× less); the speedup widened with thread count, reaching 4.2× at 32 threads (Supplementary Figure S6, left). For feature smoothing, a Rust port of `smooth_features.py` that preserves the original algorithm reduced wall-clock from 1.73 min (Python, 16 threads) to 0.61 min (Rust, single-threaded) and peak memory from 10.4 to 0.45 GiB (∼23 × less); the single-threaded Rust implementation was faster than the Python implementation at every thread count tested (Supplementary Figure S6, right). Combined, these upgrades (KaryoScope-HKS) reduce end-to-end per-haplotype wall-clock on the `repeat` feature set from 6.14 to 2.36 min on 16 threads (2.6×). Running KaryoScope with all six feature sets in a single pass adds *<*1% overhead over a single feature set (267.2 vs 264.7 s at 16 threads, KMC backend), so the runtime improvements measured here translate directly to the full pipeline output.

**Fig. S6.**
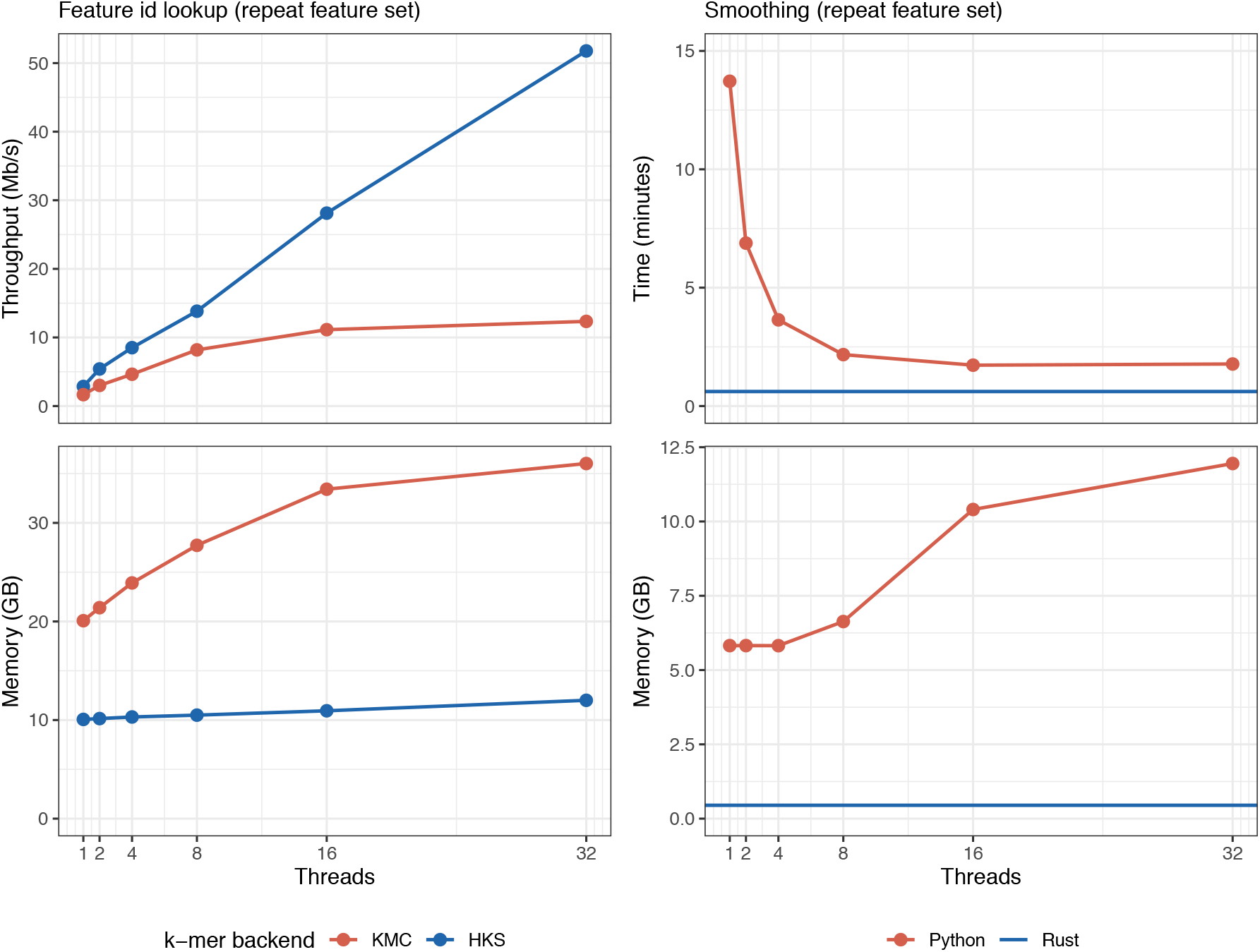
Thread scaling and upgrade path for the two principal KaryoScope annotation pipeline steps. Performance (top row) and peak resident memory (bottom row) versus thread count for *k*-mer-based feature assignment (**left column**) and hierarchical feature smoothing (**right column**) on HG002 hap1 (2.95 Gb) against the KS_human_CHM13_v2 database, measured on pinned Intel Xeon Ice Lake 2.80 GHz nodes of the TGen Gemini cluster (`repeat` feature set throughout). **Left:** Feature assignment with the released KaryoScope-KMC backend (red) vs the KaryoScope-HKS backend (blue), reported as throughput in Mb/s. HKS achieves higher throughput and lower memory at every thread count tested, with the gap widening at higher thread counts. **Right:** Smoothing with the released Python implementation (red, scaled across 1–32 threads) and a single-threaded Rust port (blue, horizontal reference), reported as wall-clock time. The single-threaded Rust implementation is faster than the Python implementation at every thread count tested and uses ∼23× less memory at 16 threads (0.45 vs 10.4 GiB).

**Fig. S7.**
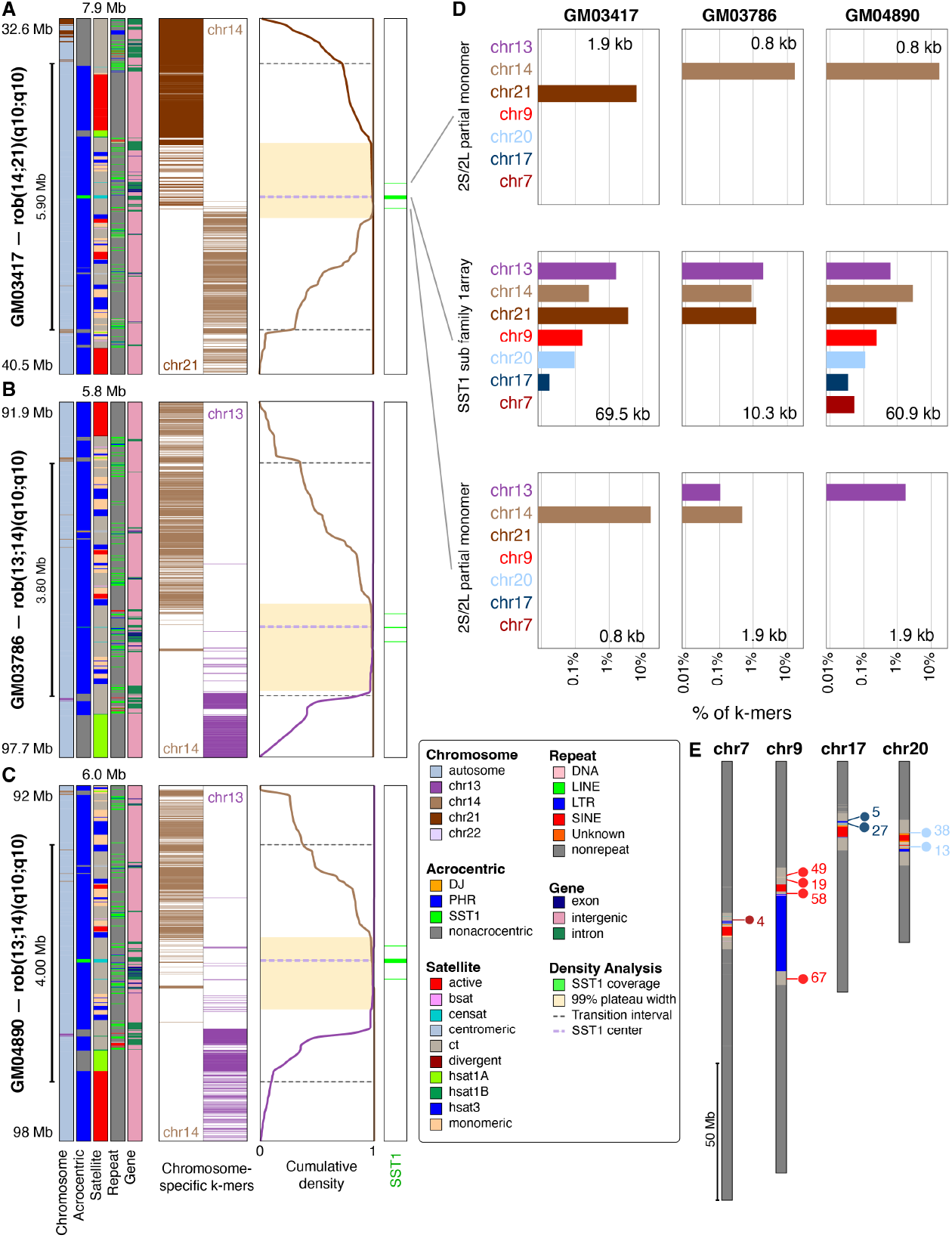
Breakpoint refinement and *k*-mer composition analysis of Robertsonian translocations. **(A–C)** Cumulative *k*-mer analysis for each fusion scaffold: **(A)** GM03417, **(B)** GM03786, **(C)** GM04890. Each panel contains four columns. The first column shows KaryoScope feature tracks (chromosome, acrocentric, satellite, repeat, gene boundaries) at the breakpoint region. The second column displays chromosome-specific *k*-mers coloured by chromosome at 100 bp resolution. The third column shows the Gaussian-smoothed cumulative curves for each fusion partner’s chromosome-specific *k*-mers (coloured by chromosome) with the 99% plateau width (yellow band), chromosome-assignment transition interval (gray dashed lines), and SST1 center (purple dashed line); the peak of the curve indicates the inferred breakpoint position. The fourth column displays SST1-specific *k*-mers (green) at 100 bp resolution. **(D)** Chromosome-specific *k*-mer composition within each SST1 array, arranged as a 3×3 grid: rows correspond to array type along the fusion scaffold (top, upstream partial array, ∼0.8–1.9 kb; middle, main array, 10.3–69.5 kb; bottom, downstream partial array, ∼0.8–1.9 kb) and columns correspond to sample (left, GM03417; middle, GM03786; right, GM04890). The main arrays correspond to the SST1 subfamily 1 (sf1) array and the flanking partial arrays correspond to the SST1 type 2L (chr13/chr21-derived) and type 2S (chr14-derived) partial monomers. Within each cell, horizontal bars show the percentage of 31-mers classified as specific to each chromosome (log_10_ scale). Bar colours follow the KaryoScope chromosome colour palette. The main arrays contain *k*-mers from the two fusion partners as well as minor contributions from non-acrocentric chromosomes (chr7, chr9, chr17, chr20). The flanking partial arrays carry *k*-mers predominantly from a single fusion partner, consistent with their position on either side of the fusion. **(E)** Positions on T2T-CHM13v2.0 of the chr7-, chr9-, chr17-, and chr20-specific 31-mers detected within the main SST1 arrays of the three ROB samples. Each chromosome is shown vertically, coloured by the KaryoScope satellite feature set. Lollipop markers indicate clusters of these 31-mers along each chromosome; numbers indicate the count of distinct 31-mers per cluster. All clusters localize to pericentromeric regions, consistent with the known pericentromeric distribution of SST1 and SST1-related elements on non-acrocentric chromosomes.

**Fig. S8.**
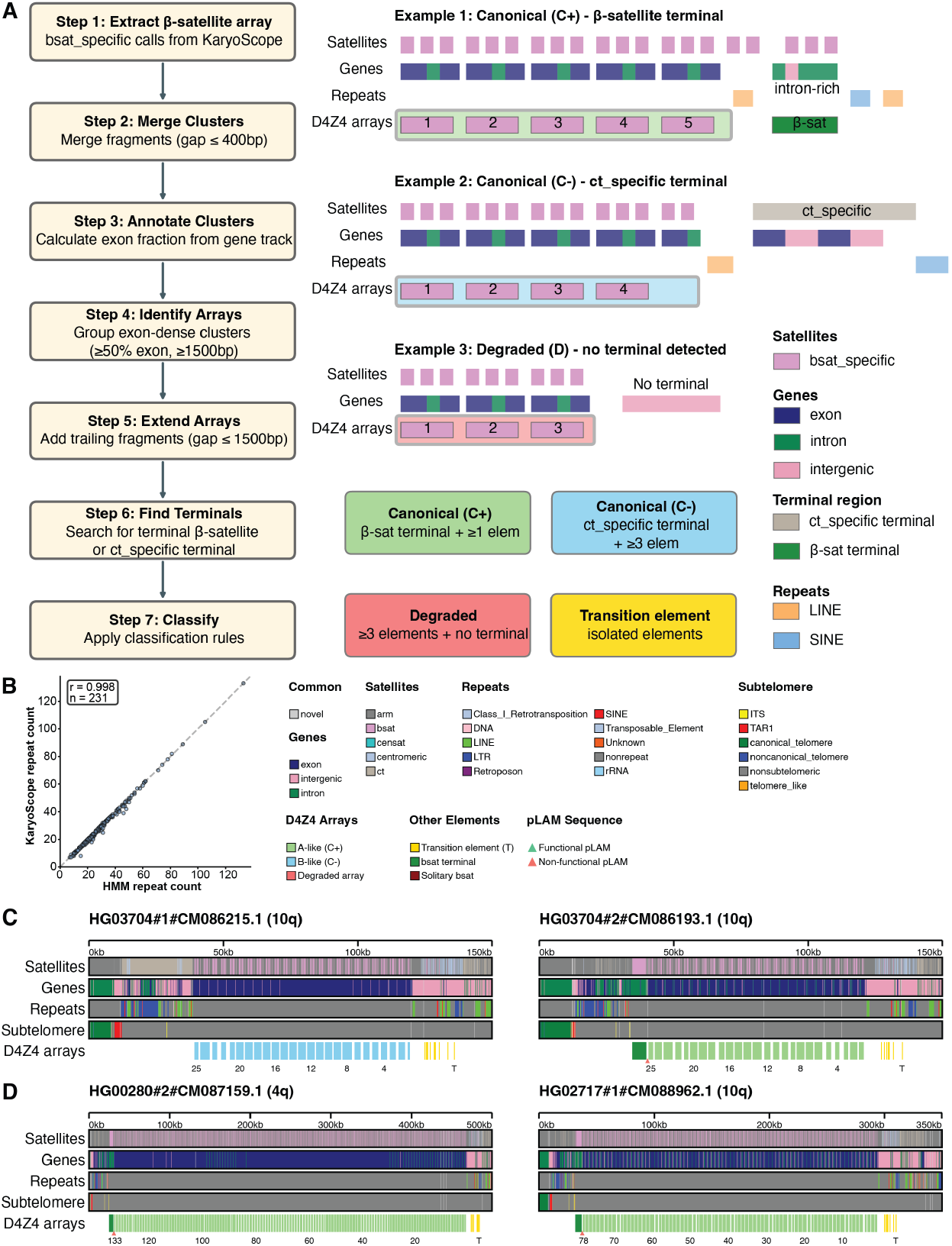
D4Z4 array classification pipeline and concordance. **(A)** D4Z4 array classification pipeline schematic. *β* -satellite fragments separated by ≤400 bp are merged into *β* -satellite repeat (BSR) clusters, clusters are annotated with exon/intron/intergenic fractions from the gene feature set, and arrays are identified as contiguous stretches of exon-dense clusters (each cluster ≥50% exon, ≥1,500 bp). Arrays are classified by their distal terminal feature as canonical C+ (A-type, *β* -satellite terminal, a pLAM proxy), canonical C− (B-type, segmental-duplication enriched terminal, ≥3 elements), or degraded (no terminal detected). **(B)** Concordance of KaryoScope-derived repeat unit counts with HMM-based counts (Pearson *r* = 0.998, *n* = 231). Dashed line indicates identity (*y* = *x*). **(C)** Additional example of a multi-array haplotype carrying two or more distinct D4Z4 tracts separated by non-D4Z4 sequence (sample HG03704, both haplotypes shown; compare Figure 4C–E). **(D)** Two haplotypes carrying exceptionally long single D4Z4 arrays: HG00280 hap2 (chr4, A-type) and HG02717 hap1 (chr10, A-type). These haplotypes are among the outliers in the haplotype asymmetry analysis (compare Figure 4H).

**Fig. S9.**
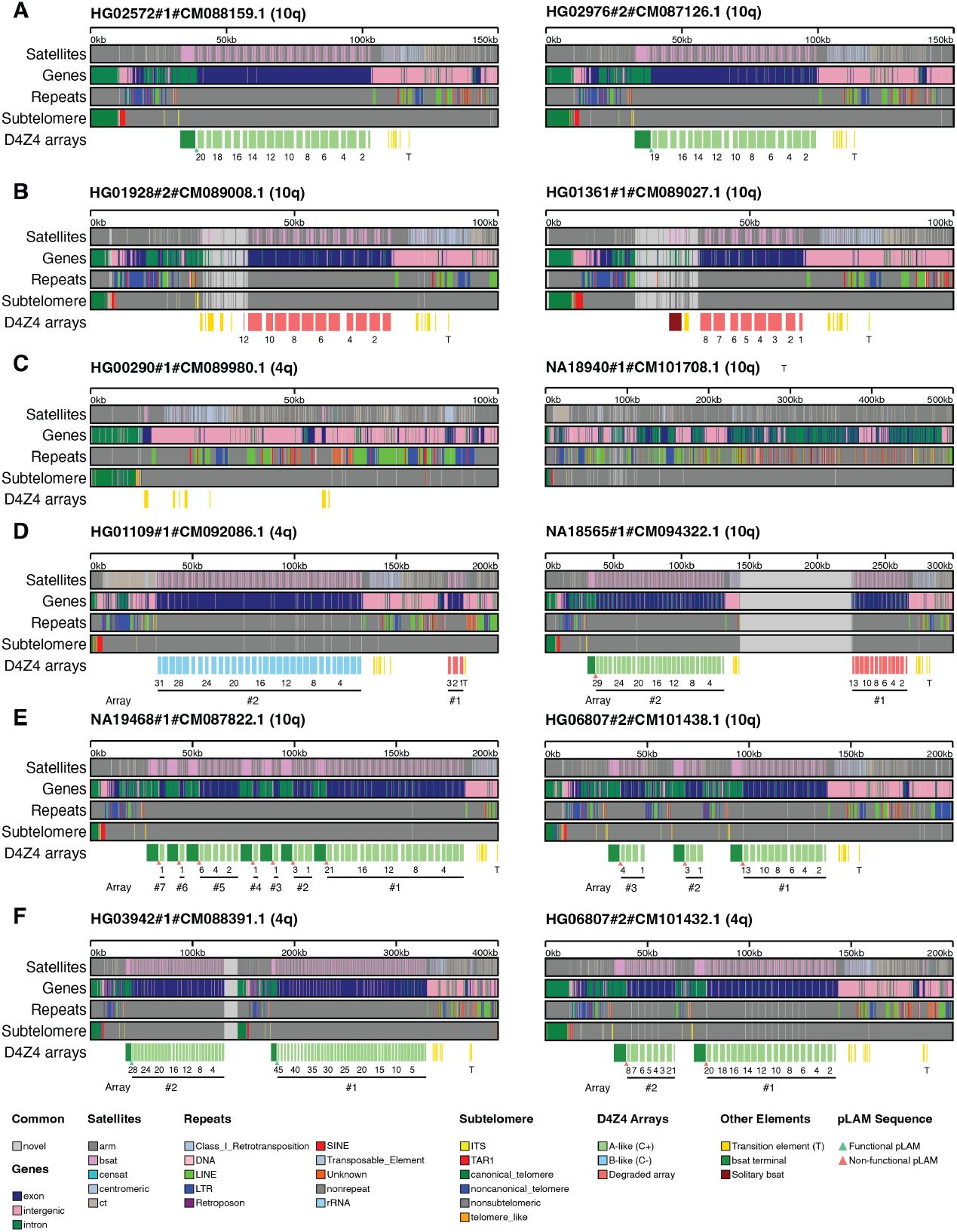
Additional examples of non-canonical D4Z4 configurations. Each panel displays satellite, gene, repeat, and subtelomeric tracks with classified D4Z4 arrays and numbered repeat units. **(A)** Chromosome 10 A-type arrays carrying functional pLAM (ATTAAA polyadenylation signal): HG02572 and HG02976. **(B)** Degraded arrays lacking canonical terminal features: HG01928 (chr10) and HG01361 (chr10). **(C)** Haplotypes with no detectable D4Z4 array: HG00290 (chr4) and NA18940 (chr10). **(D)** Multi-array haplotypes carrying two or more distinct D4Z4 tracts separated by *>*10 kb: HG01109 (chr4) and NA18565 (chr10). **(E)** Additional multi-array haplotypes on chromosome 10: NA19468 and HG06807. **(F)** Additional multi-array haplotypes on chromosome 4: HG03942 and HG06807. Note that HG06807 carries multi-array configurations on both chromosome 4 (panel F) and chromosome 10 (panel E). Compare Figure 4C–E for main-text examples.

**Fig. S10.**
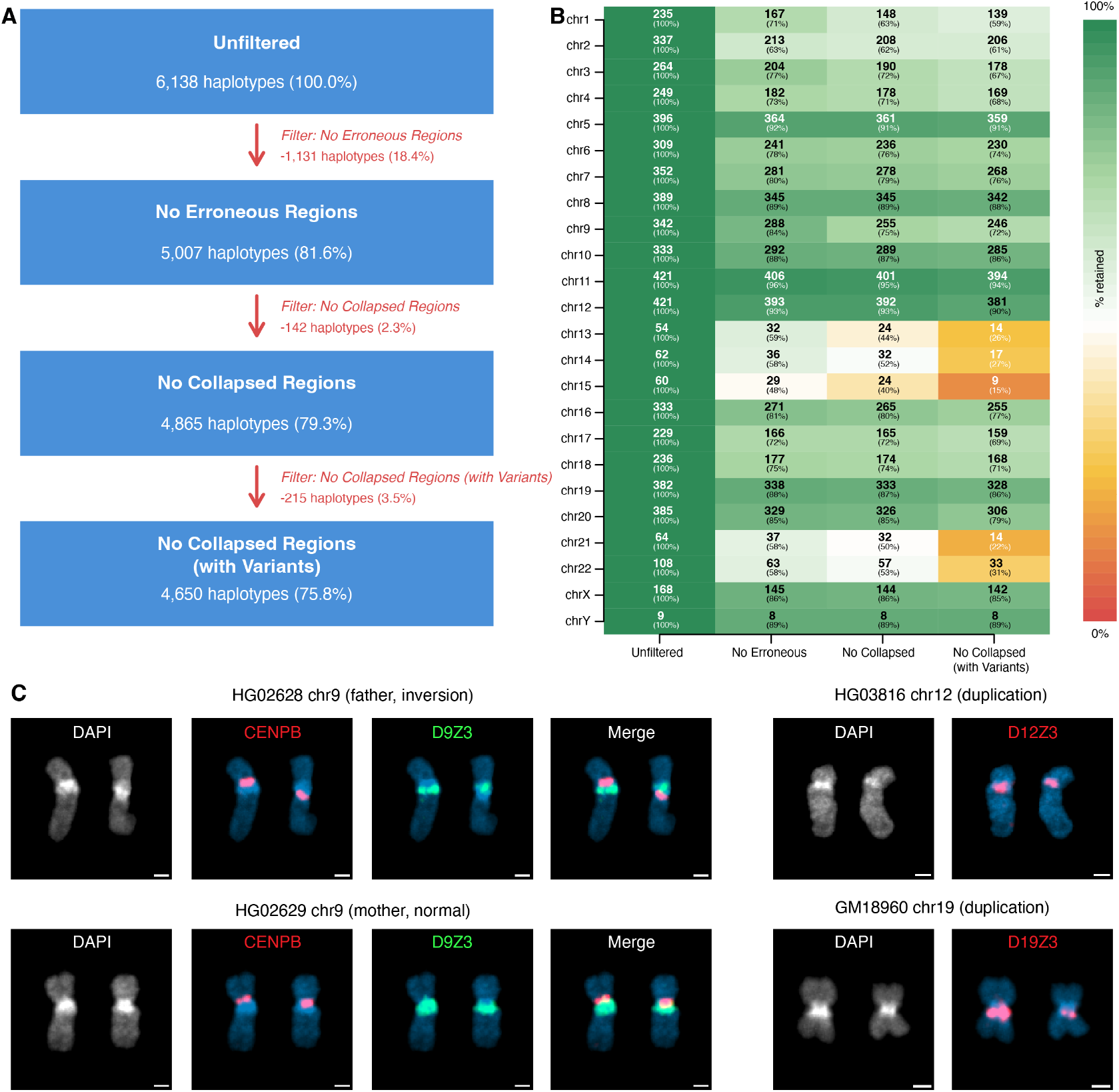
Quality filtering and additional FISH validations for the HPRC centromere census. **(A)** NucFlag QC cascade applied to the HPRC Release 2 centromere dataset. Haplotypes were sequentially filtered to remove those containing erroneous assembly regions, collapsed regions, and collapsed regions further flagged by variant calling. The retained set (4,650 haplotypes) was used as the input for Figure 5. **(B)** Per-chromosome heatmap of haplotype counts retained after the unfiltered input and each of the three filtering stages shown in panel A (columns) for chr1–chrY (rows). Cell colour encodes the percentage retained relative to the unfiltered input (scale, right). **(C)** FISH of additional minor compositional subtypes identified in Figure 5A, each performed on a cell line carrying the corresponding satellite subtype. chr9 parental FISH: father (HG02628, GWD) and mother (HG02629, GWD) probed with CENPB (red) and D9Z3 (green). HG03816 (BEB) chr12 probed with D12Z3 (red). GM18960 (JPT) chr19 probed with D19Z3 (red). DAPI counterstain and merged images are shown where available.

## Notes

https://s3-us-west-2.amazonaws.com/human-pangenomics/index.html?prefix=submissions/B5FC8EB1-5B2A-49FE-8421-6D938943DFC9--TGen_HPRCv2_KaryoScope/

https://doi.org/10.5281/zenodo.17873677

### Competing Interest Statement

The authors have declared no competing interest.

